# Structural characterization the LlaI anti-phage defense system reveals insights into the evolution of nucleotide specificity and the organization of DNA binding in McrBC restriction complexes

**DOI:** 10.64898/2026.08.31.748284

**Authors:** Anthony Q. Bui, Christopher J. Hosford, Yiming Niu, Emerson Santiago, David Moraga, Mateusz M. Wagner, Joshua S. Chappie

**Affiliations:** Department of Molecular Medicine, Cornell University, Ithaca, NY, 14853, USA; USDA-ARS Soil Management and Sugarbeet Research Unit, Crops Research Laboratory, Fort Collins, CO, USA

## Abstract

Canonical McrBC enzymes are nucleotide-powered, motor-driven endonucleases that bind and cleave modified bacteriophage DNA. Non-canonical McrBC homologs like LlaI and BsuMI are distinguished by a unique three-gene organization and the ability to target DNA site-specifically. Here, we report the atomic-resolution crystal structures of the DNA-binding module LlaI.R1 and AAA+ motor LlaI.R2 from the *Lactococcus lactis* LlaI anti-phage defense system. The crystallized LlaI.R2 hexamer traps two distinct active site conformations that correlate to different states of the nucleotide hydrolysis cycle and reveal that the organization of the critical catalytic machinery present in canonical McrB homologs is also conserved in non-canonical R2 proteins. Although canonical McrB homologs are strictly GTP-specific, we find that the R2 proteins from LlaI and BsuMI do not discriminate between different nucleotides, even when in complex with their respective R1 partners. Using mutagenesis, we define surfaces on the LlaI.R1 structure that are critical for DNA-binding and interaction with LlaI.R2. These observations support computational modelling of the assembled LlaI restriction system bound to DNA. Together, our data provide new insights into the evolution of nucleotide specificity in McrBC restriction complexes and the molecular mechanisms governing McrBC-catalyzed DNA translocation and cleavage.

**GRAPHICAL ABSTRACT:** 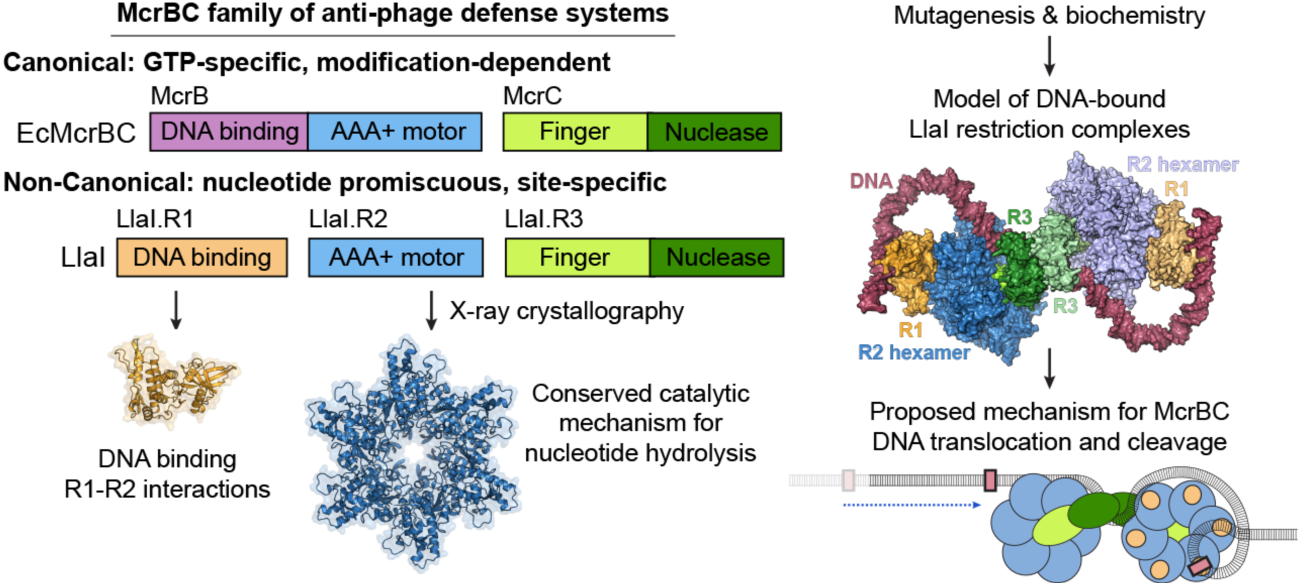

## INTRODUCTION

Bacteria have evolved a diverse collection of defense systems that destroy foreign DNA and thwart bacteriophage virus infection (1, 2). This arsenal poses a significant barrier to phage-based therapeutics and includes a plethora of different nucleases (3), which differ markedly in their specificity, structural organization, and regulation. Restriction-modification, Dnd, Ssp, BREX, and DISARM systems all bind and cleave DNA site-specifically and contain accessory components that protect the host’s genetic material by means of methylation or phosphorothioation (4–11). Modification-dependent restriction systems (MDRs), in contrast, target modified DNA (12, 13), recognizing specific patterns of cytosine methylation or glycosylation that are normally present in a subset of phage genomes (14–22). CRISPR-Cas systems function in an adaptive manner, with specificity guided by archived spacers composed of short viral DNA segments acquired from previous phage infections that direct the interference machinery to attack complementary sequences during subsequent infections (23–25). The Wadjet system guards against plasmid transformation by sensing DNA topology in closed circular DNA (26) while OLD family nucleases use ATP binding and hydrolysis to control the conformational dynamics and interactions with accessory factors that are necessary for defense functions (27–35). Structural and molecular characterization of various anti-phage systems has yielded transformative tools for molecular cloning and gene editing (5, 36–38), underscoring the general importance of these machineries to biology and medicine.

McrBC is a two-component, motor-driven endonuclease that was first identified in *E. coli* genetic screens by its ability to restrict glucosylation-deficient mutants of T4 phage (39). The *mcrB* and *mcrC* genes, which respectively encode the motor and cleavage activities of the McrBC complex, always lie in tandem within a host genome (Supplementary Figure S1A), typically localized to regions conferring host immunity (40). Canonical McrB homologs contain an N-terminal DNA binding domain (15, 17, 18, 41) and a C-terminal GTP-specific AAA+ domain that facilitates nucleotide-dependent oligomerization into asymmetric hexameric rings (42–45) (Supplementary Figure S1B). The fold and specificity of each N-terminal DNA binding domain vary between homologs, suggesting species-specific adaptations have occurred in response to different evolutionary pressures (17). *Escherichia coli* (Ec) McrB’s N-terminal SRA domain, for example, recognizes methylcytosine modifications prevalent in T even phages (13–15, 19, 46) while *Themococcus gammatolerans* (Tg) McrB’s N-terminal YTH domain preferentially binds 6-methyladenosine modifications that are present in thermophilic archaeal viruses (18). The AAA+ domain contains all the necessary catalytic machinery required for nucleotide hydrolysis, including the P loop, Walker B, and Sensor II/arginine finger motifs (45, 47). A conserved McrB consensus loop (MNT[A/S]D[R/K]S) replaces the AAA+ Sensor I motif and helps position the catalytic water for nucleophilic attack on the γ-phosphate while an aromatic residue immediately following the P loop provides a parallel π-stacking interaction that stabilizes the bound nucleotide from below (45). Canonical McrB homologs form specific hydrogen bonds with the 1’, 2’, and 6’ positions of the guanine base to recognize GTP preferentially (45). The determinants of this specificity localize to the flexible linker segment that connects the DNA binding domain to the N-terminus of the AAA+ domain (N-linker region, Supplementary Figure S1B).

McrC partners minimally consist of an N-terminal finger domain and a C-terminal PD-(D/E)XK nuclease domain (Supplementary Figure S1B). McrC cannot stably bind DNA on its own and thus must associate with the McrB hexamer to facilitate efficient DNA cleavage and restriction (44, 45, 48). In this context, McrC binding stimulates McrB’s intrinsic GTP hydrolysis ∼40-fold by repositioning the McrB consensus loop into a configuration that promotes optimal catalytic turnover (45, 49). The structural asymmetry of the McrBC complex, however, constrains McrC such that it only engages a single active site within the McrB hexamer at any given time and rotates clockwise to the next subunit to promote sequential GTP hydrolysis in a highly coordinated fashion (44, 45). *In vitro* reconstitution of Ec McrBC suggests a model for cleavage wherein complexes assemble at two R^m^C recognition sites (with R as a purine base and ^m^C as a methylated cytosine) separated by up to 3 kb and translocate DNA in a manner that depends on stimulated GTP hydrolysis (50–52). Subsequent collision of two McrBC complexes has been proposed to trigger DNA cleavage on both strands (14, 53). Cryo-EM studies have shown that Ec and Tg McrBC can dimerize through the McrC nuclease domains to form tetradecamers and that the McrC finger domain occludes the central pore of the McrB hexamer in these assemblies, preventing the potential passage of DNA through this channel (44, 45). These observations suggest that McrBC restriction complexes organize and translocate DNA by an unknown mechanism that is distinct from other hexameric AAA+ helicases and translocases (54, 55).

Our previous evolutionary analysis identified several non-canonical McrBC systems like LlaI and BsuMI that are distinguished by a unique three gene organization for the restriction machinery and the presence of accompanying methyltransferases (18) (Supplementary Figure S1A). The LlaI system resides on pTR2030, a 46.2 kb conjugal plasmid naturally found in *Lactococcus lactis* (56), and encodes a controller protein (LlaI.C), a methyltransferase (LlaI.M), a putative DNA-binding protein (LlaI.R1), an McrB-like AAA+ protein (LlaI.R2), and an McrC-like PD-(D/E)XK endonuclease (LlaI.R3) (57–59). The BsuMI system is comprised of two separate operons native to *Bacillus subtilis*: one encodes two methyltransferases (BsuMI.M1 and BsuMI.M2) while the other encodes a putative DNA-binding protein (BsuMI.R1), an McrB-like AAA+ protein (BsuMI.R2), and an McrC-like PD-(D/E)XK endonuclease (BsuMI.R3) (60, 61). Phage studies and PacBio single molecule, real-time sequencing indicate that BsuMI targets DNA site-specifically, recognizing the consensus sequence CTCGARB, where R is a purine and B is either a cytidine or a guanosine (62, 63). LlaI.M is required for efficient transformation and recovery of plasmids harboring LlaI.R1/R2/R3 together (64) and confers resistance to restriction when incorporated into a phage genome (57), suggesting the LlaI system similarly acts on DNA site-specifically (though its target sequence has yet to be identified).

To date, the architecture and molecular mechanisms of non-canonical McrBC systems remain uncharacterized. The key catalytic residues responsible for GTP hydrolysis in canonical McrB enzymes are highly conserved at the sequence level in R2 homologs (Supplementary Figure S2). The N-linker region, however, is absent in non-canonical systems as the DNA binding and motor functions are carried out by separate proteins rather than a single module built from fused domains (Supplementary Figure 1B). It is unclear how these differences affect the assembly, regulation, and functional properties of non-canonical McrBC restriction complexes.

Here, we report the atomic-resolution crystal structures of the DNA-binding module LlaI.R1 and AAA+ motor LlaI.R2 from the *Lactococcus lactis* LlaI anti-phage defense system. The crystallized LlaI.R2 hexamer traps two distinct active site conformations that correspond to either a tight, activated state primed for nucleotide hydrolysis or a post-hydrolysis open configuration that would permit nucleotide exchange. Structural comparisons with canonical McrB homologs indicate that the critical catalytic machinery required for nucleotide hydrolysis is spatially conserved in the LlaI.R2 active site. We find, however, that non-canonical R2 proteins are promiscuous with respect to nucleotide recognition, able to oligomerize with and hydrolyze GTP, ATP, XTP, and ITP. We further demonstrate that purified R1 proteins can bind both nucleotide-free R2 monomers as well as assembled R2 hexamers, and that these associations have no significant effect on nucleotide selectivity or hydrolysis. Using mutagenesis, we define surfaces on the LlaI.R1 structure that are critical for DNA-binding and interaction with LlaI.R2. These observations support computational modelling of the assembled LlaI restriction system bound to DNA. Together, our data provide new insights into the evolution of nucleotide specificity in McrBC restriction complexes and the molecular mechanisms governing McrBC-catalyzed DNA translocation and cleavage.

## MATERIAL AND METHODS

### Cloning, expression, and purification of BsuMI.R2

DNA encoding the *Bacillus subtilis* BsuMI.R2 protein (UniProt O34885) was codon optimized for *E. coli* expression and synthesized commercially by Integrated DNA Technologies (IDT), Inc. The full-length wildtype BsuMI.R2 construct (residues 1-343) was amplified by PCR and cloned into pET21b, introducing a non-cleavable 6xHis tag at the C-terminus. BsuMI.R2 was transformed into BL21(DE3) cells, grown at 37°C in 2 L of Terrific Broth to an OD_600_ of 1.0, and then induced with 0.3 mM IPTG overnight at 19°C. The cells were harvested and washed twice with nickel loading buffer (20 mM HEPES pH 7.5, 500 mM NaCl, 30 mM Imidazole, 5% glycerol (v/v), and 5 mM β-mercaptoethanol). Pellets were typically flash frozen in liquid nitrogen and stored at -80°C.

Thawed pellets from 500 mL cultures were resuspended in 30 mL of a nickel loading buffer supplemented with 10 mM PMSF, 5 mg DNase I (Roche), 5 mM MgCl_2_ and a complete protease inhibitor cocktail tablet (Roche). Lysozyme was added to 1 mg/mL and the mixture was incubated for 15 minutes with rocking at 4°C. Cells were disrupted by sonication and the lysate was cleared of debris by centrifugation at 19,685 x g for 30 minutes at 4°C. The supernatant was filtered, loaded onto a 5-ml HiTrap chelating column charged with NiSO_4_ and then washed with a nickel loading buffer. BsuMI.R2 was eluted with an imidazole gradient from 30 mM to 1 M. Peak fractions were pooled and the sample was dialyzed overnight at 4°C against HGE_50_ buffer (20mM HEPES pH 7.5, 5% glycerol (v/v), 1 mM EDTA, 1 mM DTT, and 50 mM NaCl). The sample was applied to a 5-mL HiTrap SP HP column (GE Healthcare), equilibrated with HGE_50_ buffer and then washed with HGE_50_ buffer. BsuMI.R2 was eluted with a NaCl gradient from 50 mM to 1M. Peak fractions were pooled, concentrated, and then further purified by size exclusion chromatography (SEC) using a Superdex 200 10/300 column (Cytiva), during which the protein was exchanged into a SEC_150_ buffer (20 mM HEPES pH 7.5, 150mM KCl, 5 mM MgCl_2_, and 1 mM DTT. BsuMI.R2 was concentrated to ∼30-60 mg/ml. All point mutations were introduced by QuikChange (Agilent) and the mutant proteins were purified as described for the wildtype protein.

### Cloning, expression, and purification of native LlaI.R2

DNA encoding the *Lactococcus lactis* LlaI.R2 protein (UniProt Q48593) was codon optimized for *E. coli* expression and synthesized commercially by Integrated DNA Technologies (IDT), Inc. The full-length LlaI.R2 construct (residues 1-336) was cloned via Gibson assembly into pET15bP, a modified pET15b (Novagen) plasmid in which an HRV3C protease site (LEVLFQGP) replaces the thrombin site after the N-terminal His_6_ tag. Native LlaI.R2 was transformed into BL21(DE3) cells, grown at 37°C in Terrific Broth to an OD_600_ of 1.5, and then induced with 0.3 mM IPTG overnight at 19°C. Cells were harvested and washed twice with nickel loading buffer. Pellets were typically flash frozen in liquid nitrogen and stored at −80 °C.

Thawed pellets from 2-L cultures were resuspended in 30 mL of nickel loading buffer supplemented with 10 mM PMSF, 5 µg/mL DNase I, 5 mM MgCl_2_, and a tablet of complete protease inhibitor cocktail. Lysozyme was added to 1 mg/mL and the mixture was incubated for 15 minutes with rocking at 4°C. Cells were disrupted by sonication and the lysate was cleared of debris by centrifugation at 19,685x g for 30 minutes at 4°C. The supernatant was filtered, loaded onto a 5-ml HiTrap chelating column charged with NiSO_4_, and then washed with a nickel load buffer. Native LlaI.R2 was eluted with an imidazole gradient from 30 mM to 1 M. Peak fractions were pooled, concentrated, and further purified using a Superdex 200 16/600 pg column exchanging into SEC_150_ buffer. N-terminally tagged native LlaI.R2 was concentrated to 20–40 mg/mL and used for biochemical assays. All point mutations were introduced by QuikChange (Agilent) and the mutant proteins were purified as described for the wildtype protein.

### Cloning, expression, and purification of SeMet LlaI.R2

DNA encoding full-length LlaI.R2 (residues 1-336) was cloned into pET21b and was expressed in minimal media using methionine auxotrophs (T7 Express Crystal Competent *E. coli*, New England Biolabs) according to manufacturer protocols to produce and Selenomethionine-labeled (SeMet) LlaI.R2. Thawed SeMet-labeled bacterial pellets from 500 mL cultures were purified as described for native LlaI.R2 with the following modifications: the SEC step was carried out using a Superdex 200 10/300 column (Cytiva) with the final SEC_150_ buffer supplemented with 5 mM DTT. C-terminally tagged SeMet LlaI.R2 was concentrated to 30-60 mg/ml and used for crystallography.

### Cloning, expression, and purification of BsuMI.R1

DNA encoding the *Bacillus subtilis* BsuMI.R1 protein (UniProt O35025) was codon optimized for *E. coli* expression and synthesized commercially by Twist Bioscience. The full-length BsuMI.R1 construct (residues 1-331) was amplified by PCR and cloned into the first multiple cloning site (MCS1) of pETDuet-1 (Novagen), downstream of T7 promoter-1 and containing a non-cleavable N-terminal 6xHis tag. BsuMI.R1 was transformed into BL21(DE3) cells and expressed as described above for BsuMI.R2.

Thawed pellets from 500 mL cultures were resuspended in 30 mL of a nickel loading buffer supplemented with 10 mM PMSF, 5 mg DNase I (Roche), 5 mM MgCl_2_ and a complete protease inhibitor cocktail tablet (Roche). Lysozyme was added to 1 mg/mL and the mixture was incubated for 15 minutes with rocking at 4°C. Cells were disrupted by sonication and the lysate was cleared of debris by centrifugation at 19,685 x g for 30 minutes at 4°C. The supernatant was filtered, loaded onto a 5-ml HiTrap chelating column charged with NiSO_4_, washed with a nickel loading buffer, and eluted with an imidazole gradient from 30 mM to 1 M. Peak fractions were pooled and dialyzed overnight at 4°C against TGE_50_ buffer (20mM Tris pH 8.0, 5% glycerol (v/v), 1 mM EDTA, 1 mM DTT, and 50 mM NaCl). The dialyzed sample was loaded onto a 5-mL HiTrap Q HP column (GE Healthcare) equilibrated with TGE_50_ buffer, washed with TGE_50_ buffer, and then eluted with a NaCl gradient from 50 mM to 1M. Peak fractions containing BsuMI.R.1 were pooled, concentrated, and further purified on a Superdex 200 10/300 column in SEC_150_ buffer. Purified BsuMI.R1 was concentrated to 20-50 mg/ml.

### Cloning, expression, and purification of LlaI.R1

DNA encoding the *Lactococcus lactis* LlaI.R1 protein (UniProt Q48592) was codon optimized for *E. coli* expression and synthesized commercially by Twist Bioscience. The full-length LlaI.R1 construct (residues 1-331) was amplified by PCR and cloned into pET15bP. Native LlaI.R1 was expressed as described above for BsuMI.R2.

Thawed pellets from 500 mL cultures were resuspended in 30 mL of a nickel loading buffer supplemented with 10 mM PMSF, 5 mg DNase I (Roche), 5 mM MgCl_2_, and a complete protease inhibitor cocktail tablet (Roche). Lysozyme was added to 1 mg/mL and the mixture was incubated for 15 minutes with rocking at 4°C. Cells were disrupted by sonication and the lysate was cleared of debris by centrifugation at 19,685 x g for 30 minutes at 4°C. The supernatant was filtered, loaded onto a 5-ml HiTrap chelating column charged with NiSO_4_ and then washed with a nickel loading buffer. LlaI.R1 was eluted with an imidazole gradient from 30 mM to 1 M. Peak fractions were pooled and HRV 3C protease 0.1-0.2 mg was added prior to dialysis overnight at 4°C against TGE_50_ buffer (20mM Tris pH 8.0, 5% glycerol (v/v), 1 mM EDTA, 1 mM DTT, and 50 mM NaCl). The cleaved and dialyzed sample was loaded onto a 5-mL HiTrap Q HP column (GE Healthcare) equilibrated with TGE_50_ buffer and then washed with TGE_50_ buffer. LlaI.R1 was eluted with a NaCl gradient from 50 mM to 1M. Peak fractions were pooled, concentrated, and further purified on a Superdex 200 10/300 column in SEC_150_ buffer. Purified LlaI.R1 was concentrated to 15-60 mg/ml. LlaI.R1 mutants were synthesized and cloned by Twist Bioscience and the mutant proteins were expressed and purified as described above for wildtype LlaI.R1.

### Analytical size exclusion chromatography (SEC)

For nucleotide-dependent oligomerization experiments, 100 μl samples of wildtype BsuMI.R2 or LlaI.R2 at 5 mg/mL were each incubated at 25°C for 20 minutes in absence or presence of non-hydrolyzable nucleotide analogs (2 mM GTPγS, ATPγS, XTPγS, or ITPγS) and then injected onto a Superdex 200 10/300 column equilibrated with 20 mM HEPES pH 7.5, 150 mM KCl, 5 mM MgCl_2_, and 1 mM DTT. For experiments analyzing R1-R2 binding, wildtype BsuMI.R1 and LlaI.R1 were also included with their respective R2 partner at a concentration of 5 mg/ml. Similar reaction conditions were used to assess the ability of LlaI.R1 mutants to bind to LlaI.R2 in the absence of nucleotides. The measured absorbance at 280 nm was plotted against the elution volume in each sample using KaleidaGraph (Syngergy Software).

### Size exclusion chromatography coupled to multiangle light scattering (SEC-MALS)

BsuMI.R2 and LlaI.R2 (each at 5 mg/ml) were incubated with different non-hydrolyzable nucleotide analogs (2 mM GTPγS, ATPγS, XTPγS, or ITPγS) and then loaded onto a Superdex 200 10/300 Increase column (GE Healthcare) equilibrated in 20 mM HEPES pH 7.5, 150 mM KCl, 5 mM MgCl_2_, and 1 mM DTT at a flow rate of 0.5 ml/min. The column was coupled to a static 18-angle light scattering detector (DAWN HELEOS-II) and a refractive index detector (Optilab T-rEX) (Wyatt Technology). Data were collected continuously every second. Molar mass was determined using the Astra VI software (Wyatt Technology). Monomeric BSA (Sigma) at 4 mg/ml was used for normalization of the light scattering detectors and data quality control.

### Crystallization, X-ray data collection, and structure determination of LlaI.R2

Purified native and SeMet LlaI.R2 (12 – 20 mg/mL) were incubated with 2.5 mM GDP, 25 mM NaF, and 2.5 mM AlCl_3_ and crystallized by sitting drop vapor diffusion in 0.2 M tri-ammonium citrate pH 6.0, 20% PEG 6000 with a drop size of 2 μL and reservoir volume of 65 μL. Crystals in the space group H3 typically appeared within 1-2 weeks at 20°C and were cryoprotected with Parabar 10312 (Hampton Research) prior to freezing in liquid nitrogen. Single-wavelength anomalous diffraction (SAD) (65) data were collected remotely on a single SeMet crystal with unit cell dimensions a = 176.82 Å, b = 176.82 Å, c = 64.71 Å and α = β = 90.00°, γ = 120.00° using the tunable NE-CAT 24-ID-C beamline at the Advanced Photon Source at the selenium edge energy at 12.663 keV (0.9791 Å) (Table S2). Data were integrated and scaled using XDS (66) and AIMLESS (67) via the NE-CAT RAPD pipeline. Three heavy atom sites were located using SHELX (68) and phasing, density modification, and initial model building was carried out using the Autobuild routines of the PHENIX package (69). The resulting model was incomplete and was thus used as a search model for molecular replacement into a high-resolution dataset collected from a native crystal with unit cell dimensions a = 177.75 Å, b = 177.75 Å, c = 65.03 Å and α = β = 90.00°, γ = 120.00°. Further model building and refinement was carried out manually in COOT(70) and PHENIX(69), respectively. The final model was refined to 1.80-Å resolution with R_work_/R_free_ = 18.85% / 20.67% (Supplementary Table S1) and included two LlaI.R2 monomers asymmetric unit (containing residues 1-286 and 288-337 and residues 1-286 and 294-337, respectively) along with one molecule of GDP, one molecule of citrate, two molecules of polyethylene glycol, one chloride ion, five ammonium ions, and 403 water molecules. The C-terminal 6xHis tag (LEHHHHHH) is ordered and visible in the citrate bound monomer.

### Crystallization, X-ray data collection, and structure determination of LlaI.R1

Native LlaI.R1 (8 mg/mL) was crystallized by sitting drop vapor diffusion in 0.1 M lithium citrate tribasic tetrahydrate and 20-26% PEG3350 supplemented with 1-5% 2-propanol in a drop size of 2 μL and reservoir volume of 650 μL. Crystals typically appeared within 1-2 weeks at 20°C and were cryoprotected with Parabar 10312 (Hampton Research) prior to freezing in liquid nitrogen. Crystals were of the space group C 1 2 1 with unit cell dimensions a = 80.805 Å, b = 40.349 Å, c = 99.098 Å and α = γ = 90.00°, β = 95.092°. Diffraction data from native crystals were collected remotely on the NE-CAT 24-ID-C beamline at the Advanced Photon Source at the selenium edge energy at 12.663 keV (0.97911 Å) (Table S1). Data were integrated and scaled using XDS (66) and AIMLESS (67) via the NE-CAT RAPD pipeline. The structure was solved by molecular replacement using the automated structure determination platform Auto-Rickshaw (71) and a set of pruned coordinates from an AlphaFold (72)-generated structure as the search model. Iterative model building and refinement was carried out manually in COOT (70) and PHENIX (69), respectively. The final model was refined to 1.52-Å resolution with R_work_/R_free_ = 18.69% / 21.57% (Supplementary Table S1) and contained a near complete copy LlaI.R1 (residues 1-329), six sodium ions, two chloride ions, one molecule of 2-propanol, and 274 waters.

### Nucleotide hydrolysis assays

NTPase activity was measured by using a colorimetric malachite green assay that monitors the amount of free phosphate released over time^9^. To measure the basal hydrolysis of LlaI.R2 in pET15bP, 8 or 4 μM LlaI.R2 were incubated with 1 mM NTP (ATP, GTP, XTP, or ITP) at 37°C in reaction buffer (20 mM HEPES pH 7.5, 150 mM KCl, 5 mM MgCl_2_, and 1 mM DTT). At time points of 0, 1, 2, 5, 10, 20, 30, 45, 60, 75 and 90 min, 20-μL aliquots were taken and quenched with 5 μL of 0.5 M EDTA, pH 8.0. To measure the NTPase activity of LlaI.R2 in the presence of LlaI.R1, the same conditions were used but 8 or 4 μM LlaI.R1 was added. For BsuMI.R2 in pET21b, 2 or 1 μM of protein were incubated with 1 mM NTP at 37°C in reaction buffer, and 20-μL aliquots were taken at 0, 0.5, 1, 1.5, 2, 3, 5, 10, 20, 30, and 45 minutes. To measure the NTPase activity of BsuMI.R2 in the presence of BsuMI.R1, the same conditions were used but 2 or 1 μM BsuMI.R1 was added. For colorimetric reactions, 150 μL of filtered malachite green solution were added to each sample and incubated for 5 min. The absorbance at 650 nm of the samples was measured with a Multiskan GO Microplate Spectrophotometer (Thermo Scientific). The amount of phosphate released was determined using a standard curve. To account for the spontaneous hydrolysis of NTP at 37 °C, a protein-free sample containing NTP and magnesium was incubated in parallel, and the measured amount of phosphate released at each time point was subtracted from the corresponding measurements of protein-containing samples. The specific activity is reported for all wildtype and mutant proteins (Supplementary Table S2). Quantified data represent the average of three independent experiments using multiple independently purified batches of protein with error bars indicating the standard deviation from the mean (n = 3, mean ± standard deviation). KaleidaGraph (Synergy Software) was used for statistical analysis and to plot the data.

### Microscale thermophoresis

MST experiments were conducted using a Monolith NT.115 instrument (NanoTemper Technologies) equipped with red and blue filters. BsuMI.R2 was incubated with the fluorescent RED-tris-NTA 2^nd^ generation dye (NanoTemper Technologies) in a 2:1 molar ratio at room temperature in the dark for 30 minutes before centrifugation at 13,000 rpm for 10 minutes at 4°C. The supernatant containing fluorescently labelled protein was then transferred to a new tube. Nonhydrolyzable nucleotide analogs (GTPγS, ATPγS, XTPγS, or ITPγS) were serially diluted 16 times and mixed with 50 nM of fluorescently labelled BsuMI.R2. Buffer conditions contained 20 mM HEPES pH 7.5, 150 mM KCl, 5 mM MgCl2, 1mM DTT and 0.05% TWEEN-20. Samples were incubated at room temperature in the dark for 30 minutes before being loaded into standard capillaries (NanoTemper Technologies), tested with 60% excitation power, medium MST power, and measured using M.O. Control software (NanoTemper Technologies). Data were analyzed using M.O. Affinity Analysis software (version 2.3, Nanotemper Technologies) to determine the normalized fluorescence (F_norm_) at each concentration. F_norm_ is calculated by dividing F_hot_ (average fluorescence value in the heated state) by F_cold_ (average fluorescence value measured in the cold state before the IR laser is turned on) and plotted as parts per thousand (%). F_norm_ values from at least three independently pipetted measurements were averaged (mean ± standard deviation) and plotted against the respective concentration of nonhydrolyzable nucleotide analog to obtain a binding isotherm for each substrate. Binding constants (K_d_) were determined by nonlinear curve fitting using KaleidaGraph (Synergy Software) (Supplementary Table S3). All experiments were carried out using multiple, independently purified batches of protein.

### Electrophoretic mobility shift assays (EMSAs)

The standard buffer for the EMSAs contained 10 mM Tris-HCl, pH 8.0, 250 mM NaCl, 1 mM MgCl_2_, and 1mM DTT. Binding was performed with purified LlaI.R1 (wildtype or mutants) at 25°C for 30 min in a 16-μl reaction mixture containing 5 ng/μl of λ-phage DNA (New England Biolabs). All λ-phage DNA was digested with BamHI and NdeI restriction endonucleases (New England Biolabs) at 37°C for 90 minutes and purified via a NucleoSpin gel and PCR clean-up kit (Machery–Nagel) prior to incubation with LlaI.R1. Following incubation, the samples were analyzed by 0.7% agarose gel in 1XTAE at 4°C and 80 V for 90 minutes. All gels were stained with SYBR Gold in 1XTAE overnight at 25°C (Thermo Fisher Scientific) and visualized using a Bio-Rad Gel DocTM EZ imager system.

## RESULTS

### The crystallized LlaI.R2 hexamer traps two distinct monomer conformations

To understand the architecture and catalytic mechanisms of non-canonical McrBC enzymes, we first purified the R2 proteins from the LlaI and BsuMI restriction systems. LlaI.R2 and BsuMI.R2 both form stable hexamers that can be isolated by SEC when incubated with GTPγS (Figure 1A and 1B, Supplementary Figures S3A and S4A). Although each readily crystallized in the presence of different guanine nucleotide analogs, suitable high resolution (<3 Å) diffraction could only be obtained with LlaI.R2 crystals grown in a condition containing PEG 6000, citrate, and the transition state mimic GDP.AlF_x_. We generated an initial, incomplete LlaI.R2 structure from these samples via selenium SAD phasing and then used this as a search model for molecular replacement into a 1.69-Å native dataset. The final model was refined to 1.80-Å resolution in the space group H3 with two molecules in the asymmetric unit and R_work_/R_free_ = 18.85% / 20.67% (Table S1).

**Figure 1.**
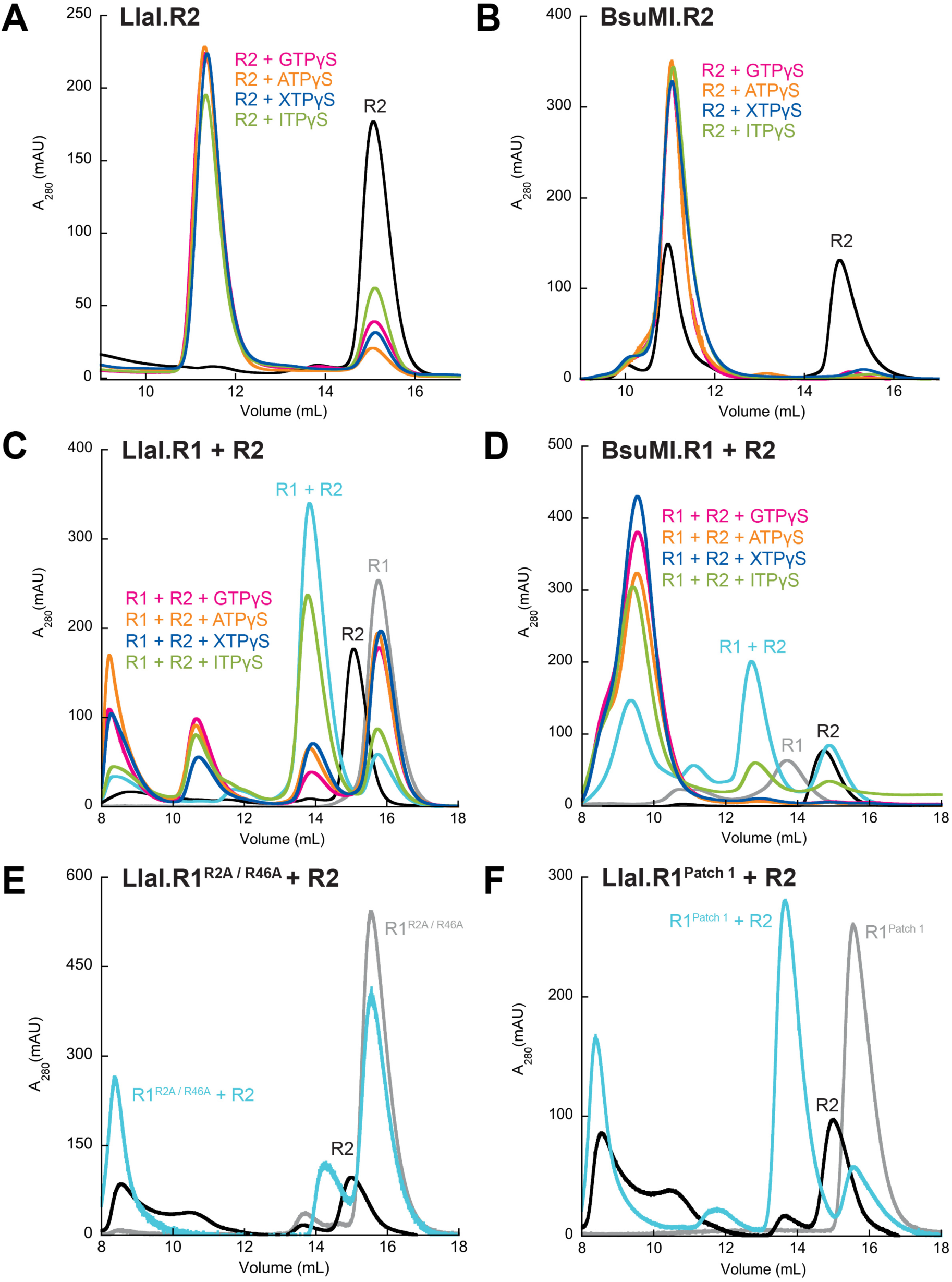
SEC analysis of nucleotide-dependent R2 protein oligomerization and R1-R2 complex formation. R2 proteins (5 mg/ml) were incubated with each nucleotide analog (2 mM) and/or their respective R1 protein partner (5 mg/ml) for 30 minutes at 25°C and then injected onto a Superdex 200 Increase 10/300 GL column. A-B. LlaI.R2 (A) and BsuMI.R2 (B) oligomerization in the absence or presence of different non-hydrolyzable nucleotide analogs (GTPγS, ATPγS, XTPγS, and ITPγS). C-D. LlaI (C) and BsuMI (D) R1-R2 complex formation and oligomerization in the absence or presence of different non-hydrolyzable nucleotide analogs (GTPγS, ATPγS, XTPγS, and ITPγS). E-F. Effects of LlaI.R1 R2A/R46A (E) and Patch 1 (R211A/R213A/K214A) mutations (F) on LlaI.R1-R2 complex formation.

Each LlaI.R2 monomer adopts a AAA+ fold with β-hairpin insertions characteristic of the H2-insert clade (73) (Supplementary Figures S5A and S6). Superpositions with the TgMcrB and EcMcrB AAA+ domains show good overall alignment of all three structures (Supplementary Figure S5B), particularly across the large subdomain. These comparisons revealed several additional structural elements that are unique to LlaI.R2, including a small helix (α2) that immediately precedes the β2 strand of AAA+ core, a helix inserted at the tip of the β6-β7 hairpin (α8), and an extra β-hairpin (β10-β11) that immediately follows β9 and packs against one end of the C-terminal helical bundle that comprises the small subdomain (Supplementary Figure S6). Structural models of BsuMI.R2 derived from AlphaFold (72), Phyre2 (74), and I-Tasser (75) also contain an additional α8 helix and β10-β11 hairpin (Supplementary Figure S6), suggesting these insertions may be an important characteristic feature of non-canonical homologs.

LlaI.R2 monomers organize within the crystal lattice as dome-shaped hexamers that pack head-to-tail in a manner resembling stacks of nestled funnels (Figure 2A, Supplementary Figure S7). The six subunits (A-F) are arranged around a central pore that spans the entire length of the hexameric assembly (Figure 2A), measuring 48 Å in diameter at the base of the dome and narrowing to 39 Å in the middle and 15 Å at the tip where the β3-β4 hairpins converge. These measurements closely match the corresponding pore dimensions in hexamers formed by the TgMcrB AAA+ domain (base diameter: 45 Å; middle diameter: 39 Å; tip diameter: 17 Å) (45), indicating that the general architecture of McrB motors is conserved even in distant homologs and across kingdoms.

**Figure 2.**
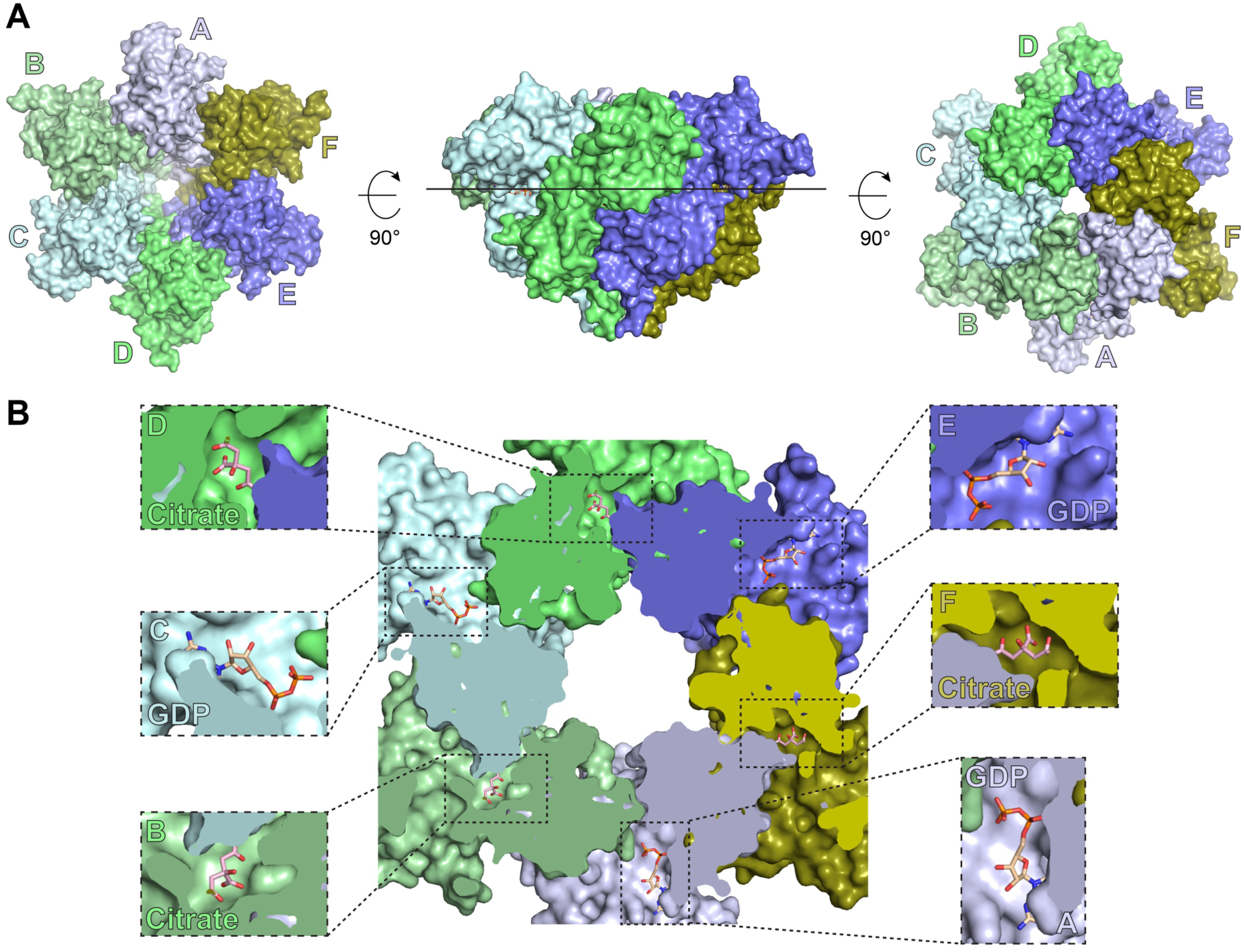
Structure of crystallized LlaI.R2 hexamer. A. Top (left), side (middle), and bottom views of the crystallized LlaI.R2 hexamer. Subunits are colored shades of blue and green and labeled sequentially “A” though “F”. B. Slice section through the LlaI.R2 hexamer at the level indicated by the black solid line shown in (A). Zoomed views highlight GDP nucleotides (subunits A, C, and E) and citrate molecules (subunits B, D, and F, derived from the crystallization condition) bound in alternating active sites.

Although LlaI.R2 crystals were grown in the presence of transition state mimic GDP.AlF_x_, we observed no electron density corresponding to the ALF_x_ moiety in any of the active sites. Instead, we found GDP and citrate bound to alternating subunits around the hexamer (Figure 2B). In subunits A, C, and E, the guanine base is stabilized through π-stacking with F235 above and R17 below while the α-and β-phosphates form hydrogen bonds with P loop main chain nitrogen atoms and the side chain of K15 (Supplementary Figure S5C). Mutation of R17 to alanine (R17A) significantly impairs LlaI.R2 oligomerization (Supplementary Figure S8A). Structurally analogous residues in other McrB homologs also form π-stacking interactions with the guanine base (44, 45) and perturbation of the corresponding W223 in TgMcrB reduces basal GTP hydrolysis (45), underscoring its critical role in stabilizing the nucleotide. Mutation of F235, in contrast, renders LlaI.R2 insoluble.

In subunits B, D, and F, citrate from the crystallization condition binds to the active site in the space normally occupied by the phosphates, where it interacts with the P loop and forms additional hydrogen bonds with main chain nitrogen atoms at the start of helix α1 (Supplementary Figure S5D). Superposition with the GDP-bound subunits shows that citrate binding induces a 21° tilt of the small subdomain (Supplementary Figure S5E), which in turn closes the nucleotide binding cleft and triggers the structural rearrangement of several active site residues (Supplementary Figure S5D). First, this movement reorients K238, allowing it to form an anchoring hydrogen bond with one of citrate’s terminal carboxyl oxygens. Second, R17 shifts upward and undergoes a rotamer flip that permits π-stacking with F235 and hydrogen bonding with Q174. In this configuration, R17 directly mimics the orientation of the guanine base, with the arginine guanidinium nitrogens spatially substituting for the nitrogens at the 1’, 2’, and 3’ positions.

Crystal packing interactions further stabilize these distinct alternating conformations within the hexamer. Closer inspection of the lattice contacts revealed that the C-terminal 6xHis tag from each citrate-bound LlaI.R2 monomer wedges into the active site of a neighboring symmetry-related GDP-bound monomer (Supplementary Figure S7A and S7B), where it forms an extensive network of hydrogen bonds that physically prevents the nucleotide sandwiching segment (NSS) from interacting with the associated GDP (Supplementary Figure S7C). The concomitant displacement of α10 forces the β-phosphate to twist away from the P loop to maintain contact with the arginine finger via non-biological hydrogen bonding interactions. We speculate that these distortions prevent the ALF_x_ from being properly stabilized along with GDP.

### The catalytic machinery and active site organization in canonical McrB proteins is conserved in non-canonical R2 homologs

GTP binds to a composite active site that is formed at the interface between adjacent subunits within the McrB hexamer, with portions of the catalytic machinery being provided both *in cis* and *in trans* (44, 45) (Figure 3 and Supplementary Figure S2). In full-length TgMcrBC and EcMcrBC restriction complexes assembled with GTPγS (PDB: 6UT5, 6UT6), the six subunits (A-F) of each McrB hexamer organize with four tight interfaces and two loose interfaces that are conformationally distinct (Figure 3A). This asymmetric arrangement directs McrC stimulation to a single active site and ensures a coordinated, sequential cycle of GTP hydrolysis around the hexamer, which is necessary for powering McrB motor functions and DNA translocation (44, 45, 52).

**Figure 3.**
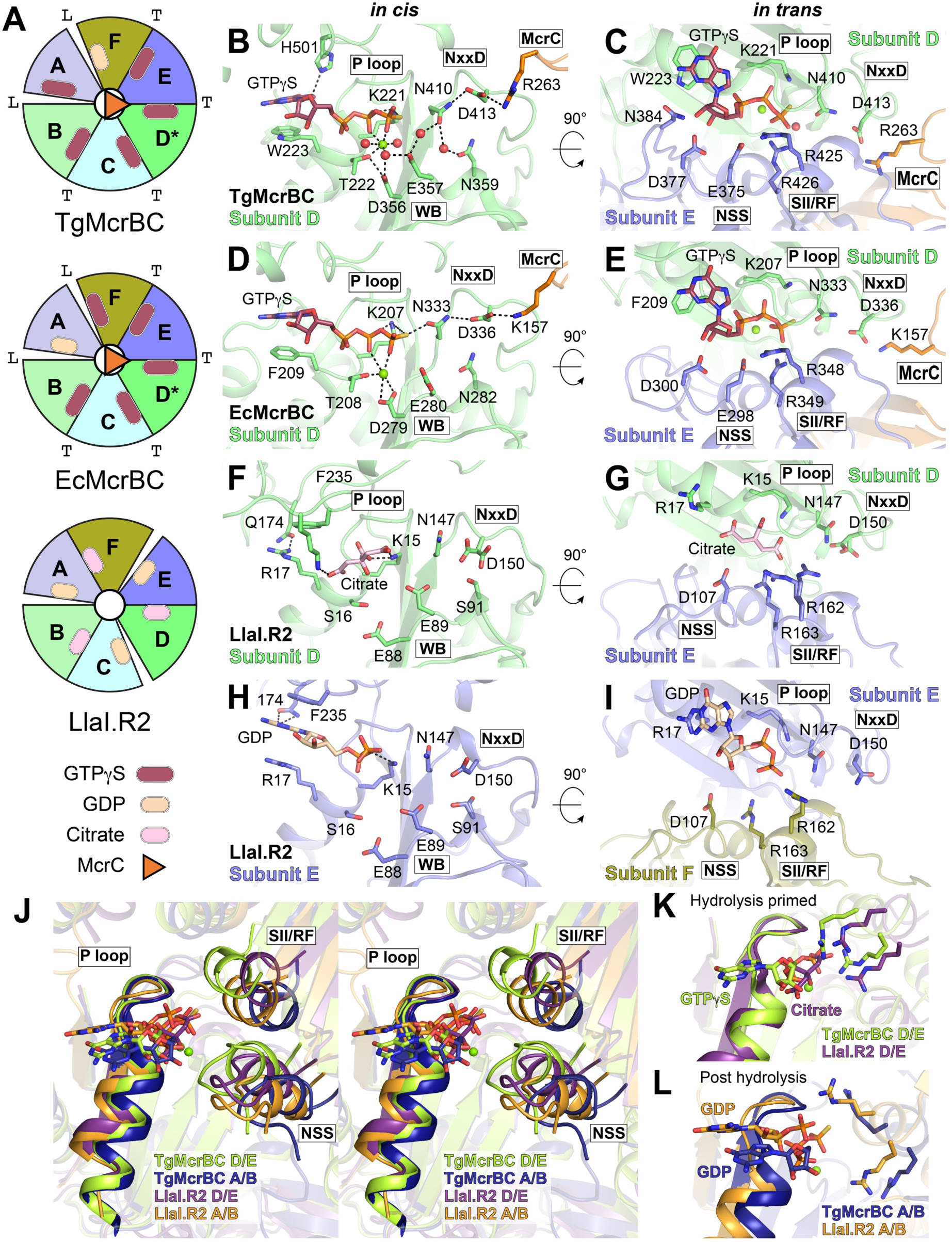
The catalytic machinery and active site organization in canonical McrB proteins is conserved in non-canonical homologs. A. Cartoon representation of subunit asymmetry in TgMcrBC (PDB: 6UT6), EcMcrBC (PDB: 6UT5), and the crystallized LlaI.R2 hexamer. McrB/R2 subunits within the hexamer are colored shades of blue and green and labeled sequentially “A” though “F”. “T” and “L” indicate the tight and loose interfaces, respectively. Asterisks denote the subunit engaged by McrC (orange triangle) for stimulated GTP hydrolysis. Colored ovals at each subunit interface represent bound GTPγS (raspberry), GDP (wheat), or citrate (pink). B-I. Zoomed views of the composite active sites formed at the TgMcrBC D/E interface (B,C), EcMcrBC D/E interface (D,E), LlaI.R2 D/E interface (F,G), and LlaI.R2 E/F interface (H,I) highlighting side chains that act *in cis* and *in trans*. Coloring and subunit labelling follow the cartoon in (A). The relative positions of the P loop, Walker B (WB), McrB consensus loop (NxxD), nucleotide sandwiching segment (NSS), sensor II/arginine finger helix (SII/RF), and McrC are labelled (see Supplementary Figure S2 for additional information). Green and red spheres denote the bound magnesium cofactor and ordered water molecules, respectively. Dashed black lines mark hydrogen bonding interactions. J. Stereo view of superposition showing the orientation of the NSS and the SII/RF helices relative to the P loop in the TgMcrB D/E interface (limon, GTPγS-bound), TgMcrB A/B interface (deep blue, GTPγS bound), LlaI.R2 D/E interface (purple, citrate-bound), LlaI.R2 A/B interface (orange, GDP-bound). K, L. Conformations of the SII/RF arginines in the citrate-bound (purple) (K) and GDP-bound (orange) (L) LlaI.R2 active sites. The corresponding positions of the SII/RF arginines in the tight TgMcB D/E interface (limon) (K) and loose TgMcB A/B interface (deep blue) (L) are superimposed for comparison.

TgMcrC engages the tight D/E interface in the TgMcrBC restriction complex (Figure 3A). Acting *in cis*, the P loop associates with the α-and β-phosphates as lysine K221 interacts with the γ-phosphate (Figure 3B). T222 coordinates the magnesium cofactor along with D356 in the Walker B motif. W223 stabilizes the guanine base from below through a π-stacking interaction while H501 hydrogen bonds with the 7’ nitrogen from above. Residues from the NSS and the α11 helix approach the nucleotide *in trans*, closing off the active site from the opposing face (Figure 3C). E375, D377, K378, and N384 form hydrogen bonds with the ribose hydroxyl groups and α-phosphate. R425 and R426 directly contact the γ-phosphate and function as the sensor II arginine and the charge-compensation arginine finger, respectively. Critically, N410 in the McrB consensus loop and E357 in the Walker B motif work together to position the catalytic water for nucleophilic attack on the γ-phosphate, thereby initiating the hydrolysis reaction (Figure 3B). TgMcrC inserts an arginine residue (R263) at the edge of this pocket and alters the conformation of the McrB consensus loop through a network of hydrogen bonds that involves N359, D413, and a bridging water molecule (Figure 3B). This action stabilizes N410 and the catalytic water in an optimal orientation, leading to a dramatic stimulation of basal GTPase activity (45). EcMcrB shares the same active site architecture and is stimulated by EcMcrC via the same conformational remodeling of the McrB consensus loop (44, 45) (Figure 3D and 3E).

Side-by-side comparisons reveal that the characteristic spatial organization of catalytic residues in canonical McrB proteins is also conserved in both the GDP-and citrate-bound LlaI.R2 monomers, with N147 poised to position the catalytic water (Figures 3F-3I and Supplementary Figure S2). An alanine substitution at this position (N147A) reduces LlaI.R2’s basal GTPase activity by 63-fold compared to the wildtype protein (Figure 4A, Supplementary Table S2), which argues that all McrB homologs hydrolyze GTP using the same underlying mechanism.

**Figure 4.**
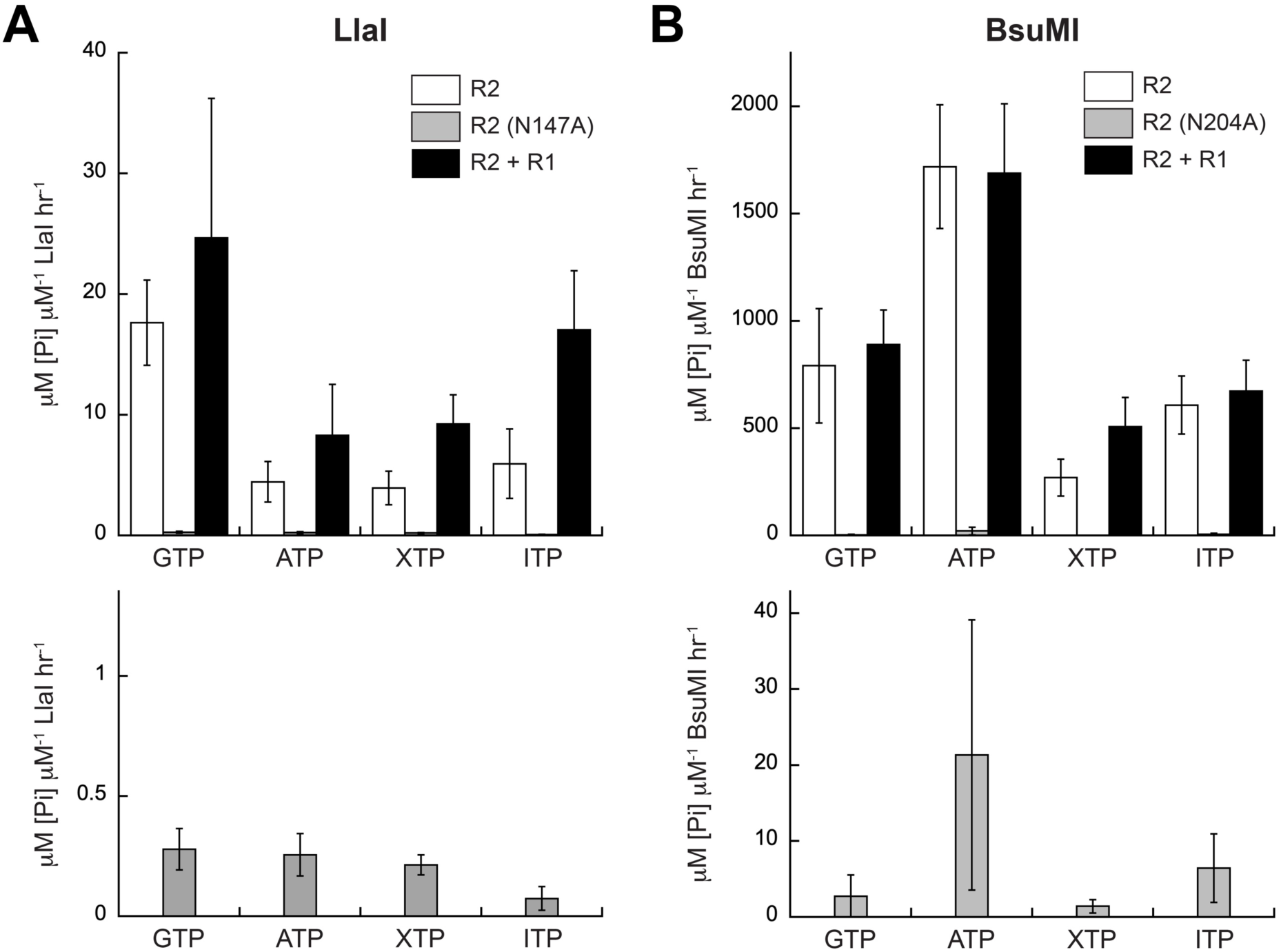
Catalytic activity of non-canonical McrB homologs. A-B. Nucleotide hydrolysis activity of wildtype LlaI (A) and BsuMI (B) R2 proteins in the absence (white) or presence (black) of their associated R1 partners. Rates were measured at 37°C with 1 mM nucleotide (GTP, ATP, XTP, or ITP) and 4 and 8 μM LlaI.R2 or 1 and 2 μM BsuMI.R2, supplemented with an equal molar amount of R1. The measured hydrolysis activities of catalytically impaired consensus loop asparagine mutants (N147A in LlaI.R2, N204A in BsuMI.R2) are included for comparison (gray), with the lower panels separately depicting these data with the Y axes rescaled for clarity. See Materials and Methods for further experimental details. Data represent the average (mean ± standard deviation) of at least three separate experiments with multiple independently purified batches of protein. Calculated specific activity values are listed in Supplementary Table S2.

Our prior cryo-EM studies demonstrated that the positions of the *trans*-acting NSS and sensor II/arginine finger (SII/RF) helices are intimately coupled to the nucleotide state and that the SII/RF arginines adopt distinct conformations throughout the GTP hydrolysis cycle (45). The NSS and SII/RF helices localize close to the nucleotide in tight TgMcrB interfaces to promote charge compensation and catalytic turnover (Figure 3J and 3K; limon) and shift away from the P loop in loose TgMcrB interfaces to permit free exchange of GDP and GTP without the need for a guanine nucleotide exchange factor (Figure 3J and 3L; deep blue). Structural superpositions indicate that the posture and positioning of the NSS and SII/RF helices in the citrate-bound LIaI.R2 active sites match the configuration of these elements found in TgMcrB tight interfaces (Figure 3J) with the SII/RF arginines directed toward the nucleotide binding pocket and primed for hydrolysis (Figure 3K). The arrangement of the NSS, SII/RF helix, and arginine side chains in the GDP-bound LlaI.R2 active sites more closely resembles a loose TgMcrB interface (Figure 3J and 3L). These observations suggest that the two unique subunit conformations captured in the crystallized LlaI.R2 hexamer represent hydrolysis-primed and post-hydrolysis states, respectively.

### R2 proteins do not discriminate between different nucleotide substrates

We previously showed that canonical McrB homologs select for GTP by forming specific hydrogen bonds with 1’, 2’, and 6’ positions of the guanine base (45). The individual structural components contributing to this readout differ in TgMcrB and EcMcrB but localize to the N-linker region, which encompasses the unstructured polypeptide segment between the fused N-terminal DNA binding domain to the start of the AAA+ α1 helix (Figure 5A and 5B, Supplementary Figures S1 and S6). Non-canonical McrB homologs lack this region as their DNA-binding R1 subunits are not tethered to their AAA+ R2 motors. As a result, we find the guanine base is left largely exposed in LlaI.R2 with nothing contacting the 1’ amine and 2’ amino group and only Q174 available to hydrogen bond with the 6’ carbonyl (Figure 5C). Mutation of this residue to alanine (Q174A) does not affect nucleotide-dependent oligomerization of LlaI.R2 (Supplementary Figure S8B), suggesting the contribution of this side chain to nucleotide binding and/or recognition is minor.

**Figure 5.**
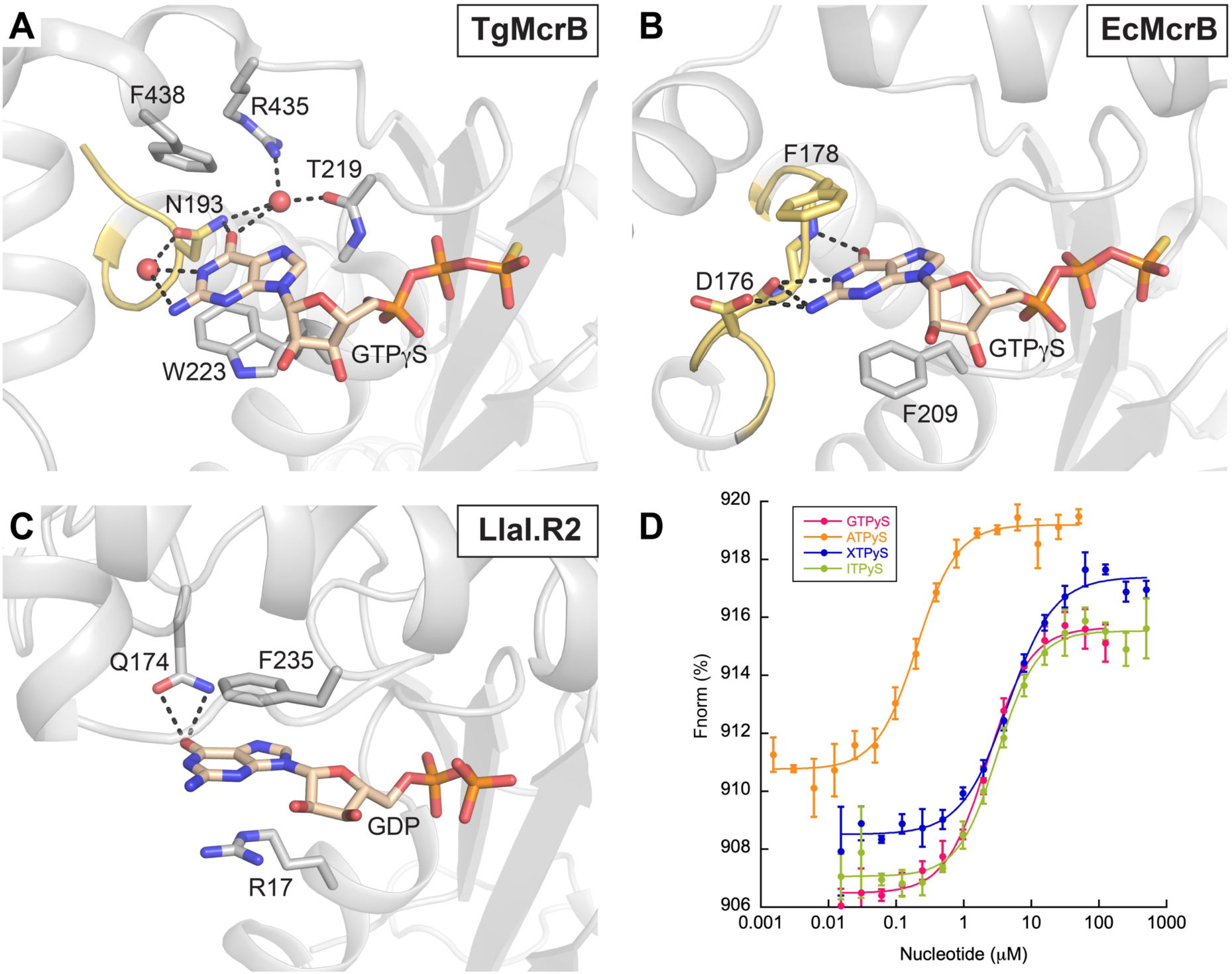
BsuMI.R2 is promiscuous for nucleotide binding. A-C. The N-linker region (yellow, see Supplementary Figures S1B and S6) containing the determinants of guanine nucleotide specificity in the canonical McrB homologs (A, *Thermococcus gammatolerans*, PDB: 6UT5; B, *Escherichia coli*, PDB: 6UT6) is absent in LlaI.R2 (C). Dashed black lines denote hydrogen bonding. Guanine nucleotides are colored wheat. Dashed black lines denote hydrogen bonds. D. MST analysis of BsuMI.R2 nucleotide-binding affinity. All data points represent average of three independent experiments (mean ± standard deviation) using multiple, independently purified batches of protein. F_norm_ is calculated by dividing F_hot_ (average fluorescence value in the heated state) by F_cold_ (average fluorescence value measured in the cold state before the IR laser is turned on) and plotted as parts per thousand (%). Binding constants were determined by nonlinearcurve fitting using Kaleidagraph (Synergy Software). Calculated K_d_ values are listed Supplementary Table S3.

Given these observations, we hypothesized R2 proteins do not discriminate between different nucleotides. To test this, we first examined nucleotide-dependent oligomerization by SEC and SEC-MALS and found that both LlaI.R2 and BsuMI.R2 can also form stable hexamers with ATPγS, XTPγS, and ITPγS (Figures 1A and 1B, Supplementary Figures S3 and S4). Prompted by this apparent promiscuity, we next asked whether the R2 proteins could also efficiently hydrolyze other nucleotide substrates aside from GTP. LlaI.R2 hydrolyzed ATP, XTP, and ITP at comparable rates, albeit about ∼3-4-fold less than GTP (Figure 4A, Supplementary Table S2). BsuMI.R2, meanwhile, showed enhanced catalytic turnover with ATP, exhibiting a ∼2-fold increase in specific activity compared to GTP and decreasing hydrolysis output with ITP and XTP, respectively (Figure 4B, Supplementary Table S2). These differences correlate with BsuMI.R2’s hierarchy of nucleotide binding as measured by microscale thermophoresis, with 11-fold higher affinity for ATPγS (Figure 5D, Supplementary Table S3). Importantly, the consensus loop asparagine mutants (N147A in LlaI.R2, N204A in BsuMI.R2) significantly impair hydrolysis activity for all nucleotides tested (Figures 4A and 4B, Supplementary Table S2). These data argue that R2 proteins hydrolyze all nucleotides using the same conserved mechanism.

### R1 proteins do not confer nucleotide specificity *in trans*

Efficient targeting of non-canonical McrBC nucleases depends on the stable association of DNA-binding R1 proteins to their R2 motor counterparts. To determine if R1 binding can influence nucleotide selectivity, we purified LlaI.R1 and BsuMI.R1 and investigated whether each could modulate nucleotide-dependent R2 oligomerization and hydrolysis activity. Using analytical SEC, we found that stable LlaI and BsuMI R1-R2 complexes form in the absence of nucleotides (Figure 1C and 1D). BsuMI.R1 and BsuMI.R2 reproducibly assembled into a large oligomeric species that elutes near the void volume and is bigger than a BsuMI.R2 hexamer when incubated together with either GTPγS, ATPγS, XTPγS, and ITPγS (Figure 1D). SEC-MALS measures the mass of the GTPγS-bound R1-R2 sample as 401297 Da, which we estimate to be an hendecamer with a stoichiometry of five BsuMI.R1 proteins bound to a BsuMI.R2 hexamer (Calculated mass: 416753 Da) (Supplementary Figure S4E). Non-hydrolyzable nucleotide analogs cause noticeable precipitation when added to LlaI.R1-R2 complexes, making it difficult to accurately assess the masses of the resulting assemblies that are formed. We speculate that this may be due to some intrinsic propensity for aggregation and/or instability that occurs with the LlaI.R1-R2 complex when it is oligomerized in the absence of the LlaI.R3 nuclease. Despite this hurdle, we note the SEC profiles produced with GTPγS, ATPγS, XTPγS, or ITPγ do not vary significantly between one another (Figure 1C). These data indicate that the formation of R1-R2 protein complexes do not depend on nucleotides and that R1 binding does not alter or impair nucleotide-dependent R2 oligomerization in any of the conditions tested.

BsuMI.R1 similarly had no significant effect on BsuMI.R2 nucleotide hydrolysis (Figure 4B, Supplementary Table S2): BsuMI.R2’s enhanced activity with ATP and distinctive pattern of catalytic output with respect to the different nucleotide substrates remained unchanged when BsuMI.R1 was included in our kinetic assays. LlaI.R1 caused a slight increase in LlaI.R2 hydrolysis with all four nucleotides (Figure 4A, Supplementary Table S2), though this change was less than three-fold even in the most extreme case with ITP. Destabilizing interface mutations that perturb McrB’s programmed asymmetry also produce a two-to three-fold stimulation of basal GTP hydrolysis as the activity of individual subunits within the hexamer become uncoupled and uncoordinated (45, 47). It is possible that LlaI.R1 binding to LlaI.R2 has a similar destabilizing effect in the absence of LlaI.R3, where the higher hydrolysis rate seen here would reflect stochastic, unfettered turnover from individual R2 monomers rather than a direct activation as occurs with McrC.

Taken together, our SEC and kinetic experiments provide indirect evidence that R1 proteins do not confer nucleotide specificity *in trans* and support the idea that non-canonical McrB homologs can utilize any nucleotide available to power the restriction nuclease complex.

### LlaI.R1 shares structural homology with eukaryotic genome maintenance proteins

Given our SEC data, we reasoned that the structural characterization of non-canonical R1-R2 complexes could provide new insights into how DNA is organized in McrBC-like restriction machines. Although structure determination of different BsuMI and LlaI R1-R2 complexes by both X-ray crystallography and cryo-EM ultimately proved unsuccessful, we were able to crystallize LlaI.R1 on its own and determine its structure at 1.52-Å resolution using a trimmed AlphaFold (72) model as a search model for molecular replacement.

LlaI.R1 crystallizes as a monomer and contains two distinctly folded domains (Figure 6, Supplementary Figure S9A). Domain 1 (residues 1-190, yellow) and is composed of a central seven-stranded β-sheet (ordered β7-β6-β4-β5-β3-β1-β2) that is flanked by clusters of α-helices. The strands are arranged in a mostly parallel fashion except for β3 and β4, which are antiparallel. Helical segments insert in loops that flank the β-sheet: helices α1 and α2 are inserted in tandem between β1 and β2; helix α3 lies between β2 and β3; helices α4 and α5 are inserted in tandem between β5 and β6; and helix α6 lies between β6 and β7. Domain 2 (residues 191-329, blue) is composed of several small groups of β-strands and α-helices. Helices α7, α8, and α9 lie in tandem at the beginning of the domain. This α-helix cluster is followed by a three-stranded β-sheet, β8-β9-β14, and two β-hairpins, β13-β12 and β11-β10. A small α-helix, α10, is sandwiched between the β-sheet and the β-hairpins. A trio of extended α-helices, α11, α12, and α13, lie in tandem towards the C-terminal end of the domain, and together with the β-sheet they form the central body of the domain. The Dali server (76) reveals that Domain 1 shares significant structural homology with the ERCC4 domains of the XPF/Mus81 family proteins Eme1 (PDB: 4P0Q, DALI1 Z score: 5.3, RMSD: 3.3 Å), ERCC1 (PDB: 6SXA, DALI Z score: 6.5, RMSD: 3.1 Å), and FAAP24 (PDB: 4BXO, DALI Z score: 6.1, RMSD: 3.0 Å) (Supplementary Figure S9B and S9C), which play critical roles in eukaryotic DNA repair processes.

**Figure 6.**
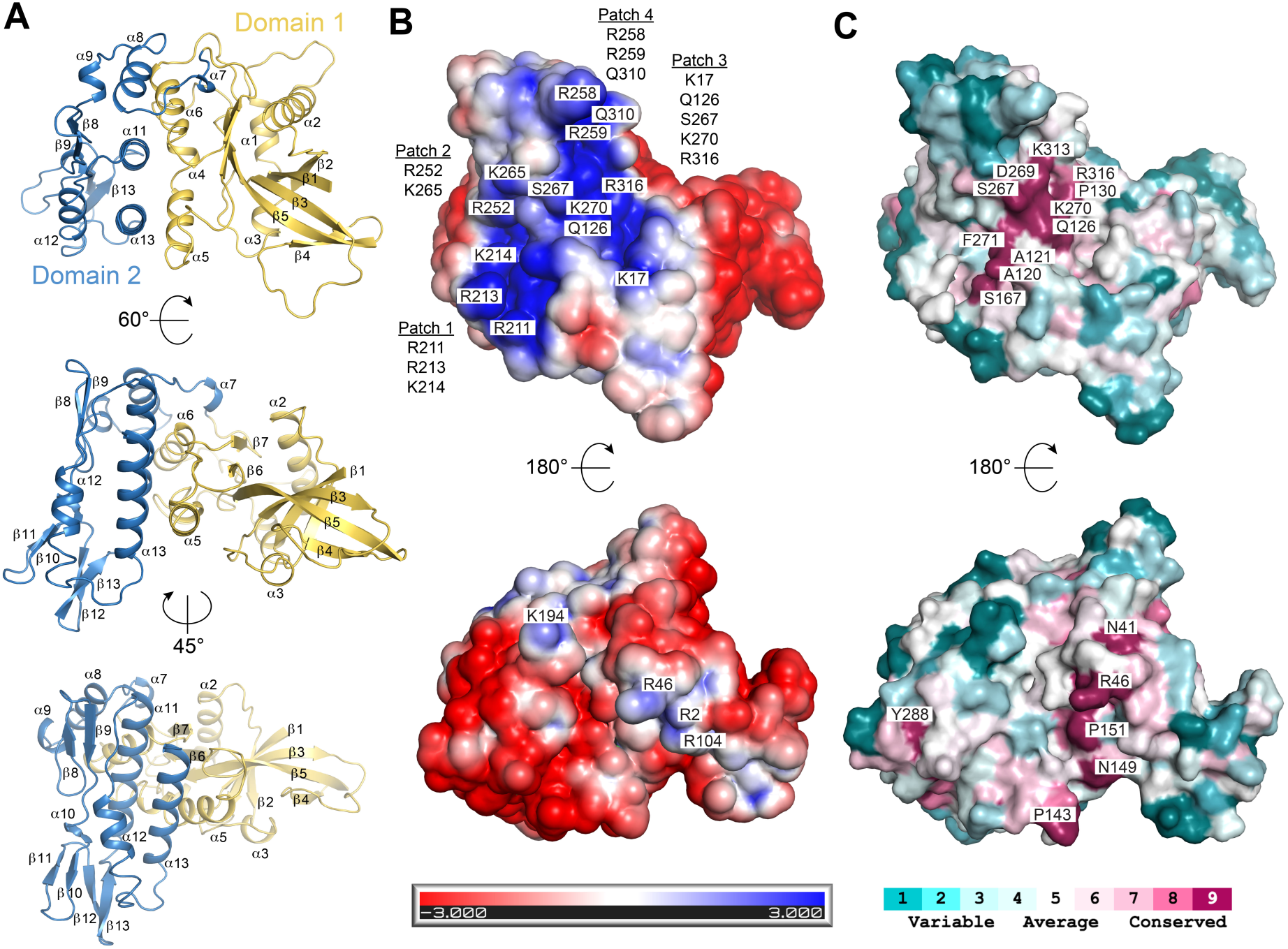
Structure of LlaI.R1. A. Crystal structure of LlaI.R1 in multiple orientations with domains 1 and 2 colored yellow and sky blue, respectively (See Supplementary Figure S9A for corresponding topology diagram). B. Electrostatic surfaces of LlaI.R1. Scale bar indicates electrostatic surface coloring from -3 K_b_T/e_c_ to +3 K_b_T/e_c_. Individual basic residues are labelled on the structure. Residues within basic patches 1-4 on the top face of the structure are enumerated separately. C. Conservation of surface exposed residues. Coloring was generated using the ConSurf server (107) and the sequence alignment in Supplementary Figure S10.

### Mutational analysis supports a model for LlaI.R1-R2 complex formation and the positioning of DNA

An extended positively charged surface comprised of four distinct basic patches (Patch 1: R211, R213, and K214; Patch 2: R252 and K265; Patch 3: K17, Q126, S267, K270, and R316; Patch 4: R258, R259, and N310) lines an extended groove that spans one face of the LlaI.R1 monomer (Figure 6B, top). The opposite face is primarily acidic, punctuated by a cluster of three arginine residues (R2, R46, and R104) and a separate lysine (K194) positioned at the distal edge (Figure 6B, bottom). ConSurf mapping indicates that neighboring residues in the immediate vicinity of these regions are highly conserved (Figure 6C, Supplementary Figure S10), suggesting that both surfaces may be functionally important.

To understand how each surface contributes to LlaI activity, we utilized AlphaFold (72, 77) to help model LlaI.R1 binding to the LlaI.R2 hexamer. We generated multiple models using different subunit stoichiometries as an unbiased starting point (1 R1:1 R2, 1 R1:2 R2, 2 R1:3 R2, 1 R1:6 R2, etc.) and found one consistent solution across all predictions that places LlaI.R1 on the exterior of the hexamer with its acidic surface contacting two R2 monomers (Supplementary Figures S11A and S11B). The putative interface buries a total surface area of ∼945 Å^2^ and includes segments of α2, the β3-β4 loop, β5, α5-β6 loop, α7, α8, α13, and β14 from LlaI.R1 and the α1-β2 loop (including α2), the α4-β5 loop, the loops flanking β6 and β7, a8, and α13 from LlaI.R2 (Supplementary Figures S6 and S12A). Key stabilizing hydrogen bond interactions are predicted between the backbone carbonyls of K327, P143, and N149 in LlaI.R1 and the Y111, S138, and T79 side chains in LlaI.R2, respectively, and between the N44 side chain in LlaI.R1 and the backbone carbonyl of S128 in LlaI.R2 (Supplementary Figure S11C). Additionally, two arginines in LlaI.R1 (R2 and R46) are predicted to form multiple anchoring contacts: the R2 sidechain hydrogen bonds with S128 in LlaI.R2 while R46 hydrogen bonds with the backbone carbonyls of Q128 and T131 and forms a salt bridge with D132 (Supplementary Figure S11C).

To validate our modelling experimentally, we first generated a series of mutations at the putative LlaI.R1-R2 interface and assessed by SEC how these changes affected LlaI.R1-R2 heterodimerization in solution. We initially tried two mutational strategies: (1) alanine substitutions that would remove stabilizing hydrogen bonds and salt bridges and (2) tryptophan substitutions that would introduce significant steric hindrance and atomic clashes. Tryptophan mutants N44W in LlaI.R1 and S128W/A130W in LlaI.R2 were both insoluble whereas the LlaI.R1 double alanine mutant R2A/R46A could be purified in milligram quantities and was found to severely impair heterodimerization with wildtype LlaI.R2 (Figure 1E). We next tested whether R2A/R46A also altered DNA binding in an electrophoretic mobility shift assay (EMSA). Although the target sequence of the LlaI restriction system is unknown, wildtype LlaI.R1 can bind digested, nonmethylated λ-phage DNA and induce a gel shift under our experimental conditions (Figure 7A) while the unrelated *B. subtilis* biosynthetic enzyme FolE cannot (Figure 7B). We hypothesized that LlaI.R1 DNA binding would remain unaffected by the R2A and R46A substitutions as these basic side chains are buried at the LlaI.R1-R2 interface in our model. As predicted, the R2A/R46A double mutant binds and shifts nonmethylated λ-phage DNA like wildtype LlaI.R1 (Figure 7C). Our biochemical observations underscore the importance of LlaI.R1 residues R2 and R46 in stabilizing the LlaI.R1-R2 interface and suggest these side chains do not contribute to DNA binding.

**Figure 7.**
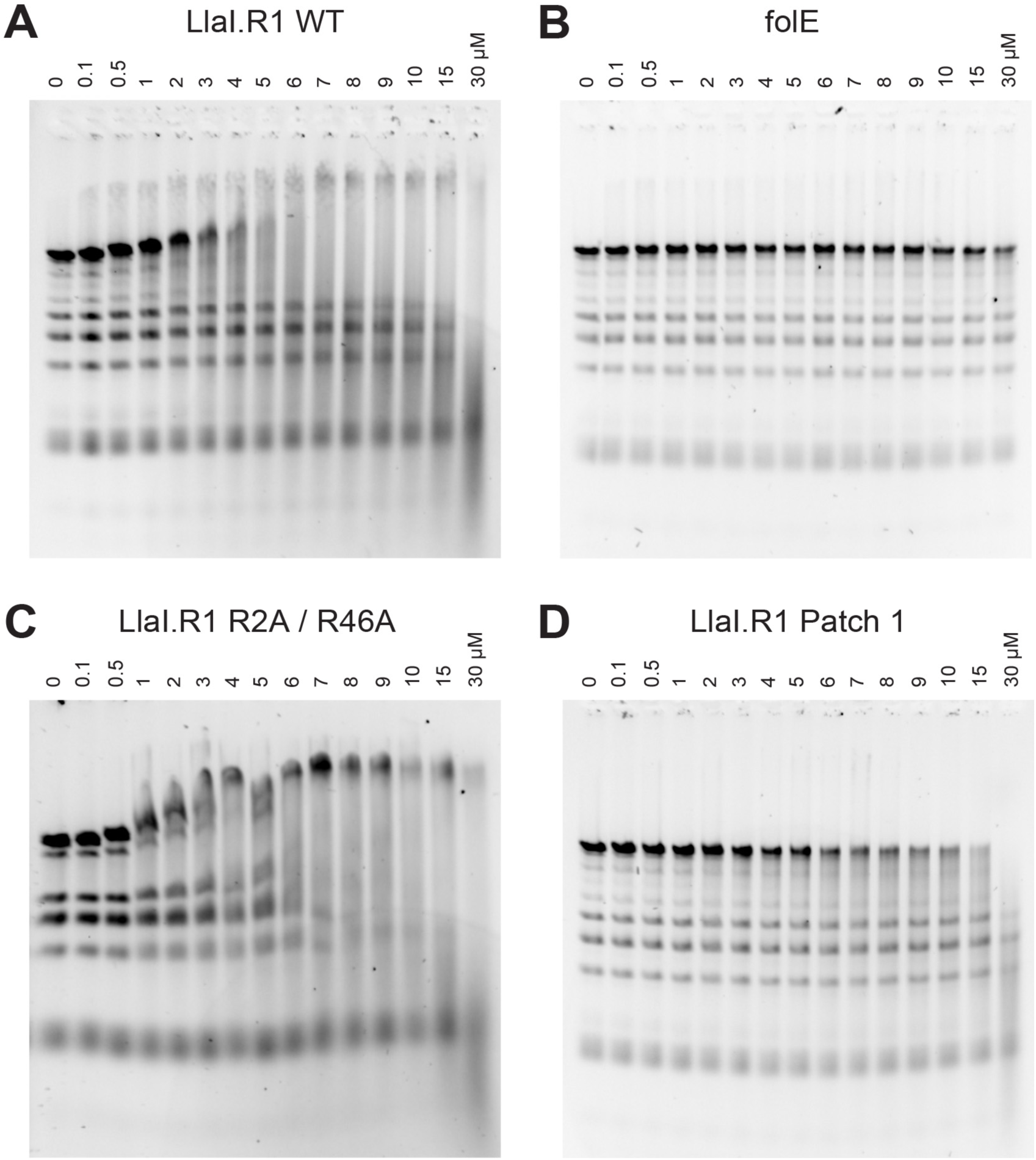
EMSA analysis of LlaI.R1 DNA binding. Samples were incubated for 30 minutes at 25°C in a 16-μL reaction mixture containing 5 ng/μL of restriction digested (BamHI/NdeI) λ-phage DNA and increasing concentrations (0 – 30 μM) of either wildtype LlaI.R1 (A), *B. subtilis* folE (B), LlaI.R1 R2A/R46A (interface-binding mutant) (C), or LlaI.R1 R211A/R213A/K214A (Patch 1 mutant) (D). Gels were stained with SYBR Green in 1X TAE buffer overnight at 25°C.

Our modelling positions LlaI.R1’s expansive positively charged surface on the exterior of the R1-R2 complex, where it would be poised to bind DNA. Computational fitting of B-form DNA (PDB: 1BNA) using different docking programs (e.g., HDOCK (78), pyDOCK (79), and NPDock (80)) supports this hypothesis, and we obtain a consistent placement the DNA duplex in an arrangement that contacts all four basic patches (Supplementary Figure S13). Mutation of Patch 1 residues to alanine (LlaI.R1 R211A/R213A/K214A) abolishes binding to nonmethylated λ-phage DNA in our EMSA assay (Figure 7D). Analogous alanine substitutions in Patches 2, 3, and 4 reduced protein solubility, hampering our ability to purify and test these mutant constructs. Importantly, however, the Patch 1 R211A/R213A/K214A triple mutant showed no defects in LlaI.R1-R2 heterodimerization (Figure 1F). Taken together, our mutagenesis data support AI-guided structural predictions and provide a credible model that details the placement of LlaI.R1 on the LlaI.R2 hexamer and illuminates the relative orientation of bound DNA within the LlaI restriction complex.

## DISCUSSION

Several defining features distinguish canonical McrB homologs from other AAA+ proteins including the presence of a nucleotide-stabilizing pi-stacking residue immediately following the P loop, a conserved McrB consensus loop replacing sensor I, and the adjacent positioning of the sensor II and arginine finger arginines on the same helix such that they act together *in trans*. Our crystallographic data confirm that these catalytic elements are not only present in LlaI.R2 but also spatially conserved within the active site (Figure 3B-3I), with N147 poised to orient the catalytic water for nucleophilic attack. AlphaFold modeling predicts a similar active site organization for BsuMI.R2 where N204 functions as the critical water-orienting side chain. Mutagenesis of these residues significantly impairs nucleotide hydrolysis in both LlaI.R2 and BsuMI.R2 (Figure 4), arguing that canonical and non-canonical McrB homologs utilize the same fundamental catalytic mechanism for basal hydrolysis. Non-canonical R2 proteins, however, bind and hydrolyze GTP, ATP, XTP, and ITP with comparable efficiency (Figure 4) and can form stable hexameric complexes with each of these nucleotides (Figure 1, Supplementary Figures S3 and S4). This intrinsic promiscuity may be advantageous in specific contexts, such as when other anti-phage defense systems that act by depleting specific nucleotide pools (e.g., dCTP and dGTP) are also present (81, 82). The R1 proteins responsible for DNA binding in non-canonical McrBC operons do not provide nucleotide selectivity *in trans* (Figure 1C and 1D), underscoring the importance of the N-linker region in conferring GTP specificity. We speculate that LlaI and BsuMI more closely resemble ancestral McrBC systems, which likely diversified over time through radical gene shuffling events driven by pressure from phages with different genomic properties. The emergence of GTP specificity in modern canonical homologs may have served to stabilize gene fusion events during this process, cementing the acquisition of new modification-dependent DNA binding domains. Evolutionary shifts away from site-specific targeting could also explain the existence of orphan methyltransferases in prokaryotes, which may persist despite being long separated from companion genes that together comprised a now defunct defense machinery.

Structural asymmetry in canonical McrBC complexes directs McrC stimulation to a single active site and dictates a coordinated, sequential cycle of GTP hydrolysis around the McrB hexamer, which has been proposed to power McrB motor functions and DNA translocation (44, 45, 52). Asymmetry is an intrinsic property of McrB proteins as evidenced by the TgMcrB hexamer showing the same pattern of tight and loose interfaces with and without McrC bound (45). LlaI.R2 crystallizes as a symmetric hexamer with two alternating active site conformations (Figures 2 and 3). This configuration arises from non-biological interactions with citrate and 6xHis tags extending from neighboring molecules in the crystal lattice (Supplementary Figures S5 and S7). While we cannot definitively rule out that non-canonical McrB homologs preferentially form symmetric assemblies, it is important to note that the citrate- and GDP-bound conformations we capture here closely mimic the “hydrolysis primed” and “post-hydrolysis” states that are present in cryo-EM reconstructions of asymmetric McrBC restriction complexes (Figure 3K and 3L). This suggests an underlying mechanistic similarity, at least with respect to catalytic activation and nucleotide exchange. It remains to be seen whether R3 protein binding induces asymmetry or simply exploits an existing feature of the R2 assembly, as is the case with McrC (45).

The pharmaceutical use of bacteriophages pre-dates the first antibiotics (83) and has gained renewed interest as more multidrug-resistant bacteria have been identified. Despite clinical success treating resistant *Acinetobacter baumannii* and *Mycobacterium abscessus* infections with phages in human patients (84, 85), anti-phage defense systems remain a significant barrier to the widespread use of phage therapy through their ability to hinder phage infection and diminish the potential for killing bacterial hosts (1, 2, 23, 86). Targeted inhibition of conserved defense machineries could therefore provide a generalized strategy to increase the efficacy of phages and broaden their use as therapeutic agents, especially as many anti-phage systems like McrBC lack eukaryotic homologs (87, 88). The LlaIR.2 crystal structure illustrates how small molecules and peptides could be employed to disrupt McrBC function. Citrate binds the LlaI.R2 active site pocket in the space normally occupied by the β and γ phosphates (Figure 3), which would prevent nucleophilic attack and catalytic turnover. 6xHis-tags associate with the NSS in the crystal lattice (Supplementary Figure 7), which blocks active site closure and would prohibit efficient nucleotide exchange. Either of these actions would uncouple the sequential, coordinated cycle of GTP hydrolysis within the McrB hexamer and ultimately disrupt the directionality of translocation. Mutations altering intersubunit coordination and/or sequential hydrolysis in other AAA+ translocases lead to loss of function and in some cases disease (89–92). These observations signal that chemical perturbations affecting McrB nucleotide binding, hydrolysis, conformational coupling, and/or symmetry would weaken host defenses thereby making drug resistant bacteria more susceptible to certain phage-based treatment regimes.

Crystallographic characterization of LlaI.R1 revealed an unexpected structural homology with the ERCC4 domains of Eme1, ERCC1, and FAAP24 (Supplementary Figure S9). While these eukaryotic genome maintenance proteins play an important scaffolding role in their respective complexes (93–96), we demonstrate that LlaI.R1 itself binds DNA (Figure 7). Docking and mutagenesis support a model where DNA is positioned across one face of LlaI.R1, contacting four distinct basic patches (Figure S13). BLAST searches across different databases identified a variety of LlaI.R1 homologs that appeared as components of different restriction system operons (Supplementary Figures S10 and S14A). AlphaFold modelling of a subset of these homologs (Supplementary Figure S14B) predicts that they all share a common two-domain fold with an extensive basic surface spanning the same face as LlaI.R1 (Supplementary Figure S15A and S15B), hinting at a conserved mode of DNA binding. Although DNA can be positioned in roughly the same orientation in each model, the distribution of side chains contacting the DNA backbone changes in each homolog (Supplementary Figure S15C). These differences likely reflect individual specificity preferences as non-canonical McrBC homologs have been evolved to bind DNA in a sequence-specific manner. Strikingly, all versions of AlphaFold fail to build a high-confidence model for BsuMI.R1 even when paired with its R2 and R3 partners, which may be a consequence of there being fewer homologs available that share true sequence homology across the entire protein. Future studies dissecting BsuMI.R1 DNA binding will be particularly informative as a complement to our findings here since its target sequence has been identified (62, 63).

Nucleic acid substrates typically pass through the central pore of hexameric AAA+ helicases and translocases (54, 55). In McrBC restriction complexes, the McrC finger domain specifically inserts into this region of the McrB hexamer, blocking the passage of DNA through the central channel (44, 45). DNA must therefore be organized in a configuration that is different from other AAA+ motors. Our ability to delineate the pathway of DNA has been hindered by the limitations of past structural models: DNA is absent in all previous McrBC cryo-EM structures and the tethered N-terminal DNA binding domains within these three-dimensional reconstructions are highly mobile and poorly resolved (44, 45), making it impossible to computationally fit corresponding DNA-bound N-terminal domain structures determined by X-ray crystallography (15, 18, 19). Our LlaI structures provide a unique opportunity to address these ambiguities and gain further insights into the organization of McrBC restriction complexes and molecular mechanisms governing their function. The R2A/R46A double mutant validates the predicted R1-R2 interface (Supplementary Figure S11) as it disrupts LlaI.R1 binding to wildtype LlaI.R2 (Figure 1E) without affecting DNA binding (Figure 7C). Leveraging this information, we constructed a model for the fully assembled LlaI restriction system bound to DNA (Supplementary Figure S16). This assembly contains two LlaI complexes – each comprised of one R1 DNA binding protein, one R2 hexamer, and one R3 nuclease – that are dimerized via the R3 PD-(D/E)xK nuclease domains in a manner that follows the organization of McrBC tetradecamers captured by cryo-EM (PDB: 6UT6, 6UT7, 6HZ5, 6HZ6, 6HZ7, 6HZ8, 6HZ9) (Supplementary Figure S16A). The R3 nuclease dimer straddles DNA such that the active sites of the opposing monomers are each oriented against a different DNA strand (Supplementary Figure S17A), thereby facilitating complete cleavage of the duplex through a metal-dependent mechanism that is shared by Type II restriction endonucleases and other PD-(D/E)xK family enzymes involved in DNA recombination and repair (8, 97, 98) and conserved across McrC homologs (48) (Supplementary Figure 17B-17D).

With the pore of each R2 hexamer occluded by the R3 finger domain (Supplementary Figure 16B), DNA would need to follow a trajectory that passes through the R3 nuclease dimer and then wraps around each LlaI complex, bending at distinct points to permit binding to the exterior R1 proteins while avoiding steric clashes with nearby R2 subunits (Supplementary Figure S16C and S16D). This arrangement, which shares aspects with some early models for McrBC assembly (99), would explain why McrBC enzymes always cleave DNA between two recognition sites (46, 50) and is consistent with observations that cleavage does not occur when recognition sites are spaced too close together (e.g., <40 bp for EcMcrBC) (50, 51, 53).

Structural superpositions show that the TgMcrB AAA+ domain has several large insertions that overlap with the putative R1-R2 interface in LlaI and would sterically clash with the positioning of LlaI.R1 (Supplementary Figure S12B). EcMcrB similarly has unique insertions in the AAA+ small subdomain that coincide with LlaI.R2’s α13 helix and significantly alter the physical contours and chemical properties of this region (Supplementary Figure S12C). These differences imply that McrB AAA+ domains may have co-evolved to accommodate the incorporation of new recognition modules, with species-specific insertions constraining how binding domains can be integrated within the core motor and cleavage machinery. This is reflected in our previous McrBC cryo-EM reconstructions where unassigned densities corresponding to the Tg and EcMcrB DNA binding domains localize to different places in each structure (45). We anticipate that the minimal spacing of recognition sites required for efficient cleavage as well as the degree of DNA bending will differ between McrBC homologs depending on how their DNA binding domains are situated in the assembled complex.

*E. coli* McrBC is a valuable diagnostic tool that has been utilized extensively for mapping methylation patterns associated with human disease (100, 101), human development (102), plant immunity (103), and genomic regulation (104, 105). The modular nature of the McrBC family suggests that its members could be readily adapted to other diagnostic applications and may serve as promising scaffolds for creating new programmable endonucleases with customized specificities, particularly given the wide assortment of nucleic acid binding domains present in different McrB homologs (18, 87). Successful engineering of novel McrBC motor fusions will likely require extensive mutagenesis within the AAA+ domain to stabilize new targeting domains along with careful design of the N-linker sequence to preserve the molecular interactions that dictate nucleotide selectivity.

Drawing from parallels with Type I and Type III restriction systems, it has been proposed that DNA translocation facilitates the convergence of two McrBC complexes such that their collision triggers DNA cleavage (44, 52, 99, 106). Based on our collective structural and biochemical findings, we propose an alternative mechanistic model that illustrates how a single McrBC tetradecamer could mediate DNA translocation and cleavage on its own, with the strain imposed on DNA by McrB conformational rearrangements serving the primary activator of McrC’s nuclease activity (Figure 8). In this scheme, heptameric McrBC restriction complexes (6 McrB:1 McrC) initially dimerize via the McrC PD-(P/E)xK nuclease domains to form stable tetradecamers in solution (Figure 8, Step 1). Binding to a single recognition site would position the tetradecamer with DNA anchored to the exterior face of one McrB hexamer and loosely threaded through the dimerizing McrC nuclease domains, making minimal interactions with the other McrB hexamer (Figure 8, Steps 2-3). Sequential rounds of McrC-stimulated GTP hydrolysis powered by concomitant nucleotide exchange (GTP for GDP) and phosphate release (Figure 8, Step 4, right) would induce counterclockwise rotation of both McrB hexamers (Figure 8, Step 4, left). This action in turn would generate DNA looping and/or twisting near the bound recognition site, resulting in directional translocation as DNA is pulled toward one side of the complex (Figure 8, Step 5). Engagement of a second recognition site by the unbound McrB hexamer would exert an opposing force on DNA (Figure 8, Step 6), creating a strained conformation that stalls directional movement and ultimately promotes McrC-catalyzed DNA cleavage (Figure 8 Step 7). Such a mechanism does not explicitly exclude collisional activation as a translocating tetradecamer would be similarly stalled if it encountered another DNA-bound McrBC complex or if other physical roadblocks that prevented further directional movement prior to engagement of the second recognition site. An important caveat is that these models are derived from structural snapshots and bulk biochemical experiments and do not account for factors like DNA topology, which has been shown to affect EcMcrBC activity *in vitro* (52). Detailed analysis of McrBC function by single molecule approaches will be necessary to clarify the underlying mechanochemical properties further and examine how DNA topology influences McrBC loading and/or biases directional movements and cleavage.

**Figure 8.**
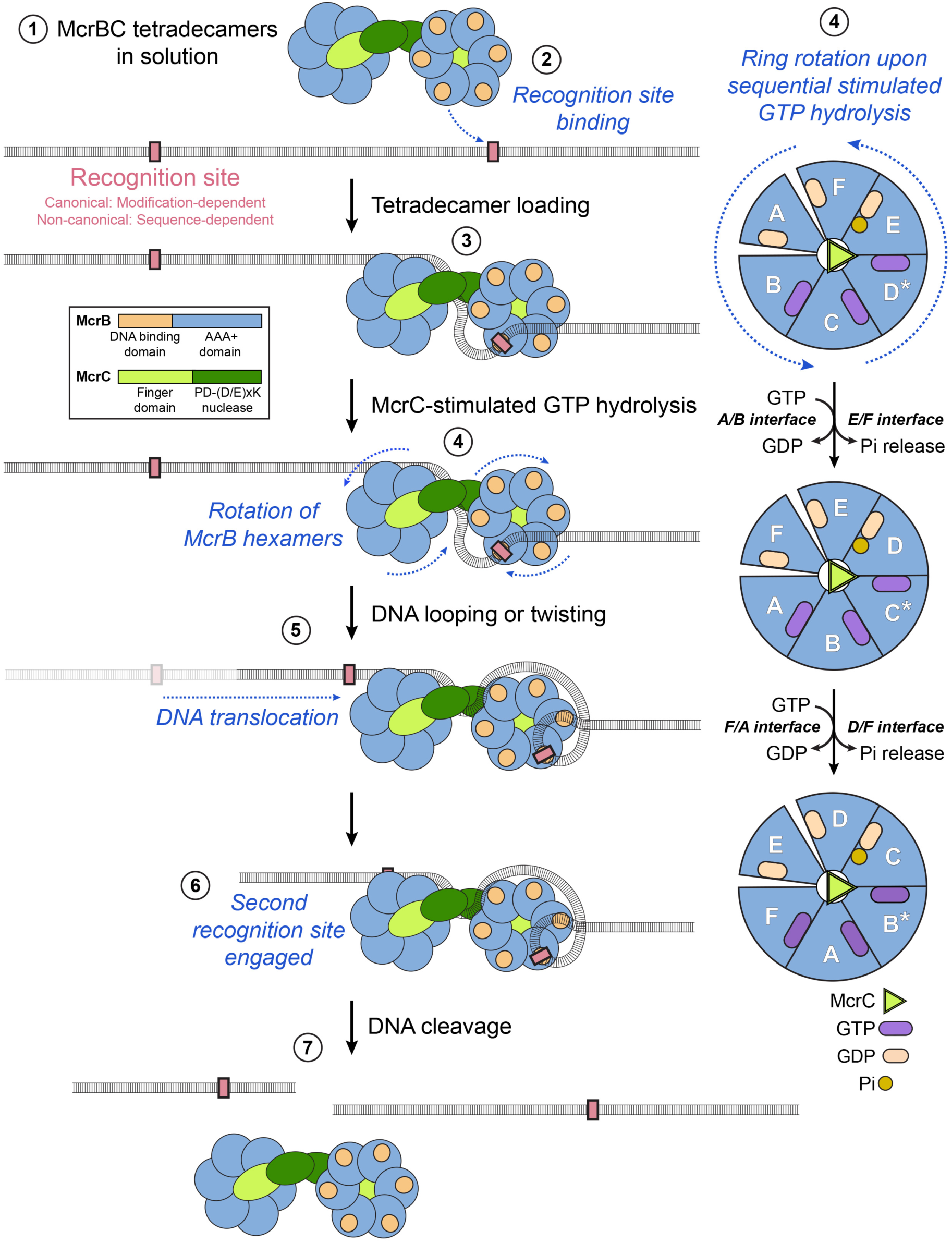
Mechanistic model for McrBC DNA translocation and cleavage. Diagram of proposed cleavage pathway for McrBC homologs (left) with accompanying schematic illustrating rotational movement of McrB hexamer following sequential rounds of McrC stimulated GTP hydrolysis (right). Heptameric McrBC restriction complexes (6 McrB:1 McrC) dimerize via the McrC PD-(P/E)xK nuclease domains to form tetradecameric assemblies in solution (Step 1). Binding to a single recognition site targets the tetradecamer to DNA (Step 2), where it loads with DNA anchored to one McrB hexamer and loosely threaded through the dimerizing nuclease domains (Step 3). Sequential rounds of McrC-stimulated GTP hydrolysis driven by concomitant nucleotide exchange (GTP for GDP) and phosphate (Pi) release cause rotation of McrB hexamers (Step 4), which causes DNA looping/twisting that facilitates translocation (Step 5). Engagement of a second recognition site by the opposing McrB hexamer (Step 6) creates a strained conformation that promotes DNA cleavage (Step 7).

## Supporting information

Bui et al. Supplementary Information

## ACKNOWLEDGEMENTS

We thank Drs. Joseph Peters, John Helmann, and Marijn Ford for insightful discussions and critical reading of the manuscript and the Northeastern Collaborative Access Team (NE-CAT) beamline staff at the Advanced Photon Source (APS) for assistance with remote X-ray data collection. We extend a special thank you to Dr. Kevin Dorn for his prowess in physically manipulating random office supplies to illustrate the potential conformational rearrangements diagrammed in our cartoons.

## AUTHOR CONTRIBUTIONS

**Anthony Q. Bui**: Conceptualization, Methodology, Investigation, Validation, Visualization, Writing – original draft, Writing – reviewing and editing. **Christopher J. Hosford**: Conceptualization, Methodology, Investigation, Validation. **Yiming Niu**: Investigation, Validation. **Emerson Santiago**: Investigation, Validation. **David Moraga**: Investigation, Validation. **Mateusz M. Wagner**: Investigation, Validation. **Joshua S. Chappie**: Conceptualization, Methodology, Investigation, Validation, Data Curation, Visualization, Resources, Funding Acquisition, Supervision, Writing – original draft, Writing – reviewing and editing.

## SUPPLEMENTARY DATA

Supplementary Data are available online.

## CONFLICT OF INTEREST

None to declare.

## FUNDING

This work was supported by the National Institutes of Health Grant GM120242 (to J.S.C.) and the USDA-ARS Appropriated Project Number 3012-21220-011-000D. The NanoTemper Monolith NT.115 instrument used for MST binding experiments was funded by an equipment supplement from the National Institutes of Health (GM120242-02S1 to J.S.C). A portion of this work is based upon research conducted at the Macromolecular Diffraction facility at the Cornell High Energy Synchrotron Source (MacCHESS) supported by the National Science Foundation [DMR1332208] and the National Institutes of Health [GM103485] and the NE-CAT beamlines (24-ID-C and 24-ID-E) supported by the National Institutes of Health [P41 GM103403, S10 RR029205]. This research also used resources of the Advanced Photon Source, a U.S. Department of Energy (DOE) Office of Science User Facility operated for the DOE Office of Science by Argonne National Laboratory under Contract No. DE-AC02-06CH11357. The USDA is an equal opportunity provider and employer. Mention of trade names or commercial products in this publication does not imply recommendation or endorsement by the USDA. Funding for open access charge: USDA-ARS Appropriated Project Number 3012-21220-011-000D.

## DATA AVAILABILITY

The atomic coordinates and structure factors for the LlaI.R2 and LlaI.R1 crystal structures have been deposited in the Protein Data Bank with accession codes 38DJ and 38DK, respectively.

## Notes

### Competing Interest Statement

The authors have declared no competing interest.

