## Supplementary material for "Structural characterization the LlaI anti-phage defense system reveals insights into the evolution of nucleotide specificity and the organization of DNA binding in McrBC restriction complexes": Bui et al. Supplementary Information

**Supplementary Tables**

Table S1. X-ray data collection and refinement statistics.

Table S2. R2 protein nucleotide hydrolysis activity.

Table S3. Dissociation constants for BsuMI.R2 nucleotide binding from MST experiments.

**Supplementary Figures**

Figure S1. Operon and domain organization of McrBC homologs. Related to Figure 1.

Figure S2. Sequence conservation of catalytic motifs required for McrB GTP hydrolysis. Related to Figures 1 and 2.

Figure S3. SEC-MALS analysis of LlaI.R2 complexes. Related to Figure 1.

Figure S4. SEC-MALS analysis of BsuMI.R2 complexes. Related to Figure 1.

Figure S5. LlaIR.2 AAA+ fold and comparison of GDP- and citrate-bound LlaI.R2 monomers. Related to Figure 2.

Figure S6. Topology diagrams of the AAA+ domains from canonical and non-canonical McrB homologs. Related to Figures 2-5.

Figure S7. LlaI.R2 crystal packing and 6xHis tag interactions. Related to Figures 1 and 2.

Figure S8. SEC analysis of nucleotide-dependent oligomerization of mutant LlaI.R2 proteins. Related to Figures 1 and 2.

Figure S9. LlaI.R1 shares structural homology with eukaryotic genome maintenance proteins. Related to Figure 6.

Figure S10. Sequence alignment of representative LlaI.R1 homologs. Related to Figures 1, 6, and 7.

Figure 11. Structural modelling of LlaI.R1-R2 interactions. Related to Figures 1, 6 and 7.

Figure S12. Evolutionary constraints affecting the association of the DNA-binding and AAA+ modules in McrBC homologs. Related to Figures 1 and 6.

Figure S13. Structural modelling of LlaI.R1 DNA binding. Related to Figures 6 and 7.

Figure S14. Structural modelling of LlaI.R1 homologs. Related to Figures 6 and 7.

Figure S15. Structural comparison of LlaI.R1 with modelled R1 homologs identifies a conserved mode of DNA binding. Related to Figures 6 and 7.

Figure S16. Structural modelling of LlaI restriction complexes bound to DNA. Related to Figures 2, 6, 7, and 8.

Figure S17. Modelled LlaI.R3 interactions with DNA. Related to Figures 2, 6, 7, 8, and Supplementary Figure S16.

**Table S1. X-ray data collection and refinement statistics.**

| | SeMet LlaI.R2 | LlaI.R2<br>PDB: 38DJ | LlaI.R1<br>PDB: 38DK |
|---|---|---|---|
| **Data collection** | | | |
| Space group | H3 | H3 | C 1 2 1 |
| Cell dimensions | | | |
| *a*, *b*, *c* (Å) | 176.82, 176.82, 64.71 | 177.75, 177.75, 65.03 | 80.805, 40.349, 99.098 |
| α, β, γ (°) | 90, 90, 120 | 90, 90, 120 | 90, 95.092, 90 |
| Resolution (Å) | 88.41 – 2.30 (2.38 – 2.30) | 88.87 – 1.69 (1.72 – 1.69) | 98.97 – 1.52 (1.55 – 1.52) |
| $R_{sym}$ or $R_{merge}$ | 15.8 (162.8) | 6.1 (324.6) | 0.089 (1.081) |
| $R_{meas}$ | 18.3 (186.5) | 6.3 (340.5) | 0.096 (1.296) |
| $CC_{1/2}$ (%) | 97.8 (39.6) | 100 (26.9) | 0.998 (0.602) |
| $I / \sigma I$ | 9.2 (0.9) | 21.8 (0.7) | 10.4 (1.0) |
| Completeness (%) | 99.8 (99.8) | 99.5 (92.2) | 98.6 (83.1) |
| Redundancy | 4.1 (4.1) | 12.7 (10.9) | 6.7 (5.8) |
| **Phasing** | | | |
| Initial F.O.M. | 0.65 | | |
| Number of sites | 3 | | |
| **Refinement** | | | |
| Resolution (Å) | | 88.87 – 1.80 | 98.71 – 1.52 |
| No. reflections | | 70917 (2799) | 47685 (6339) |
| $R_{work}$ / $R_{free}$ (%) | | 18.85 / 20.67 | 18.69 / 21.57 |
| No. atoms | | | |
| Protein | | 5413 | 2738 |
| Ligand/ion | | 71 | 12 |
| Water | | 403 | 274 |
| *B*-factors | | | |
| Protein | | 40.90 | 25.93 |
| Ligand/ion | | 50.20 | 39.79 |
| Water | | 45.25 | 34.51 |
| R.m.s deviations | | | |
| Bond lengths (Å) | | 0.009 | 0.008 |
| Bond angles (°) | | 1.237 | 1.027 |
| **Ramachandran statistics** | | | |
| Favored (%) | | 98.05 | 97.86 |
| Allowed (%) | | 1.95 | 2.14 |
| Outliers (%) | | 0 | 0 |

*Values in parentheses are for highest-resolution shell. Each dataset was derived from a single crystal.

**Table S2. R2 protein nucleotide hydrolysis activity.**

| System | Sample | Nucleotide | Specific activity | Standard deviation |
|---|---|---|---|---|
| LlaI | R2 | GTP | 17.63 | 3.53 |
| | | ATP | 4.46 | 1.69 |
| | | XTP | 3.94 | 1.39 |
| | | ITP | 5.95 | 2.87 |
| | R2 (N147A) | GTP | 0.28 | 0.086 |
| | | ATP | 0.26 | 0.088 |
| | | XTP | 0.21 | 0.041 |
| | | ITP | 0.074 | 0.049 |
| | R2 + R1 | GTP | 24.66 | 11.57 |
| | | ATP | 8.30 | 4.23 |
| | | XTP | 9.2 | 2.40 |
| | | ITP | 17.06 | 4.89 |
| BsuMI | R2 | GTP | 791.62 | 265.85 |
| | | ATP | 1719.56 | 288.42 |
| | | XTP | 269.77 | 85.76 |
| | | ITP | 607.87 | 135.29 |
| | R2 (N204A) | GTP | 0.80 | 3.48 |
| | | ATP | 21.36 | 17.78 |
| | | XTP | 1.42 | 0.86 |
| | | ITP | 6.43 | 4.51 |
| | R2 + R1 | GTP | 890.13 | 161.78 |
| | | ATP | 1688.63 | 324.054 |
| | | XTP | 507.38 | 136.72 |
| | | ITP | 673.79 | 143.75 |

Nucleotide hydrolysis was measured using a colorimetric malachite green assay that monitors the release of free phosphate (Pi) over time (1). Specific activity is defined as the amount of phosphate released per hour per micromolar of protein ($\mu M^{-1}$ Pi $\mu M^{-1}$ protein $h^{-1}$). Values represent the average of at least three separate experiments with multiple individually purified batches of protein.

**Table S3. Dissociation constants for BsuMI.R2 nucleotide binding from MST experiments.**

| Protein | Construct | Nucleotide | Kd (µM) | Error (%) |
|---|---|---|---|---|
| BsuMI.R2 | WT | GTPγS | 2.26 | 17.56 |
| | | ATPγS | 0.20 | 1.89 |
| | | XTPγS | 4.47 | 43.70 |
| | | ITPγS | 3.17 | 29.52 |

MST binding isotherms used to determine $K_d$ values are shown in Figure 5B.

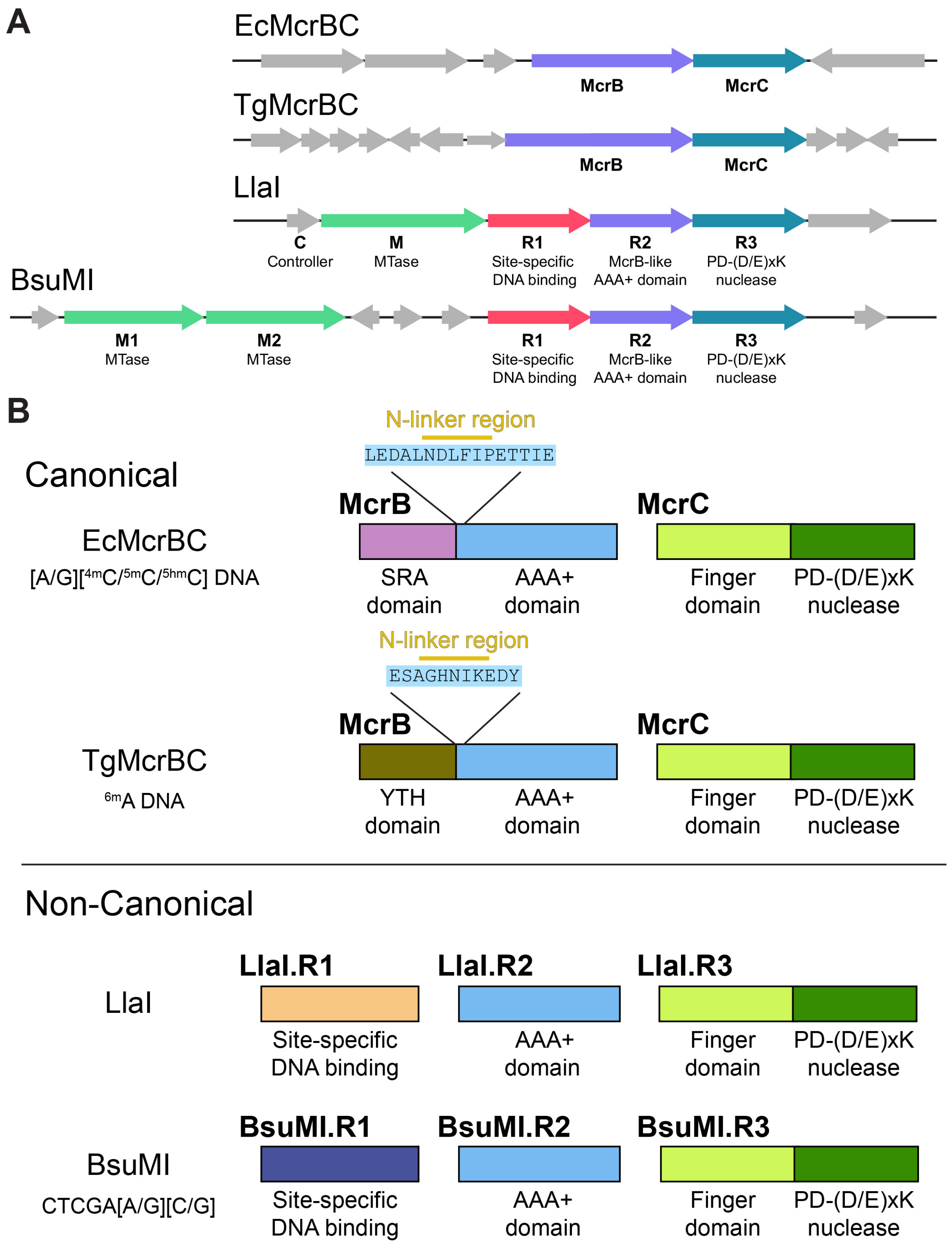


**Figure S1. Operons and domain organization of McrBC homologs.** A. Gene neighborhoods of representative McrBC restriction systems. Canonical systems from *Escherichia coli* (Ec) and *Themococcus gammatolerans* (Tg) are shown along with the non-canonical systems LlaI from *Lactococcus lactis* and BsuMI from *Bacillus subtilis*. B. Domain organization of McrBC enzymes. Sequences of the N-linker region that confers guanine nucleotide specificity in canonical homologs are shown. The recognition site targeted by each system varies. EcMcrBC binds R$^{m}$C sites, where R is a purine and $^{m}$C is 4-methyl- ($^{4m}$C), 5-methyl- ($^{5m}$C), or 5-hydroxymethylcytosine ($^{5hm}$C) (2–4). TgMcrBC binds 6-methyladenosine-modified ($^{6m}$A) DNA (5). BsuMI binds DNA site-specifically at the sequence CTCGARB, where R is a purine and B is either a cytidine or a guanosine (6, 7).

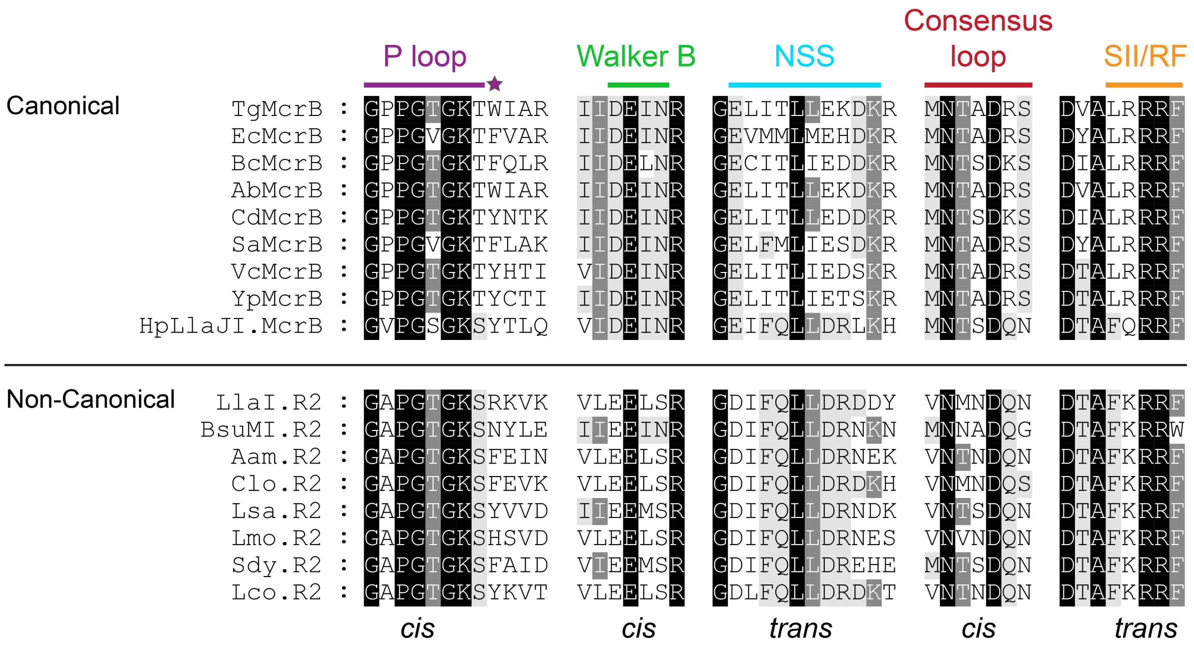


**Figure S2. Sequence conservation of catalytic motifs required for McrB GTP hydrolysis.** Representative homologs from canonical and non-canonical McrBC restriction systems are shown with shading indicating conservation: white text on black background, 100% conserved; white text on dark gray background, 70% conserved; black text on light gray background, 50% conserved. Purple star denotes the $\pi$-stacking residue that stabilizes the nucleotide base. See Supplementary Figure S7 for additional topological context. Abbreviations with the associated NCBI accession numbers or DOE IMG/M (8) IDs are as follows: Tg, *Thermococcus gammatolerans* EJ3 (644807740); Ec, *Escherichia coli* MG1655 (NP_418766); Bc, *Bacillus cereus* 03BB102 (ACO28538); Ab, *Aciduliprofundum boonei* T469 (646649070); Cd, *Clostridioides difficile* R20291 (646337923); Sa, *Staphylococcus aureus aureus* MRSA252 (637153557); Vc, (WP_229767288) Yp, *Yersinia pestis* sv. Orientalis CO-92 (637199492); HpLlaJI, Helicobacter pylori JM99 (637022177); Aam2110, *Anaerosphaera aminiphila* DSM 21120 (SHH63134); Clo, *Clostridium* sp. DSM 8431 (SFU59101); Lsa, *Ligilactobacillus salivarius* BCRC 14759 (ATP37642); Lmo, *Listeria monocytogenes* PIR00545 (AVU90227); Sdy, *Streptococcus dysgalactiae* subsp. *equisimilis* AC-2713 (CCI63035); Lco, *Loigolactobacillus coryniformis* subsp. *torquens* DSM 20004 = KCTC 3535 (ATO43927).

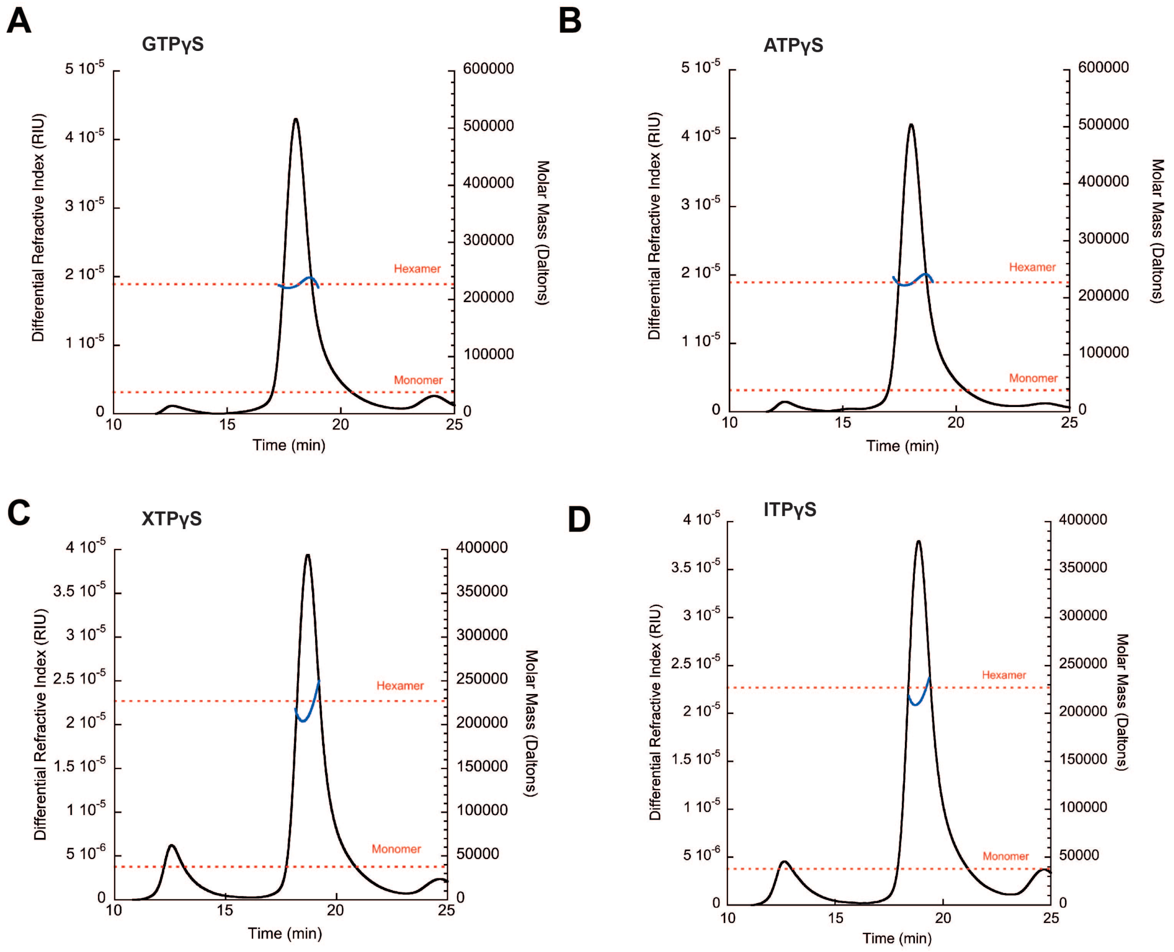


**Figure S3. SEC-MALS analysis of LlaI.R2 complexes.** A-D, SEC-MALS of LlaI.2 in the presence of GTPγS (A), ATPγS (B), XTPγS (C), and ITPγS (D).

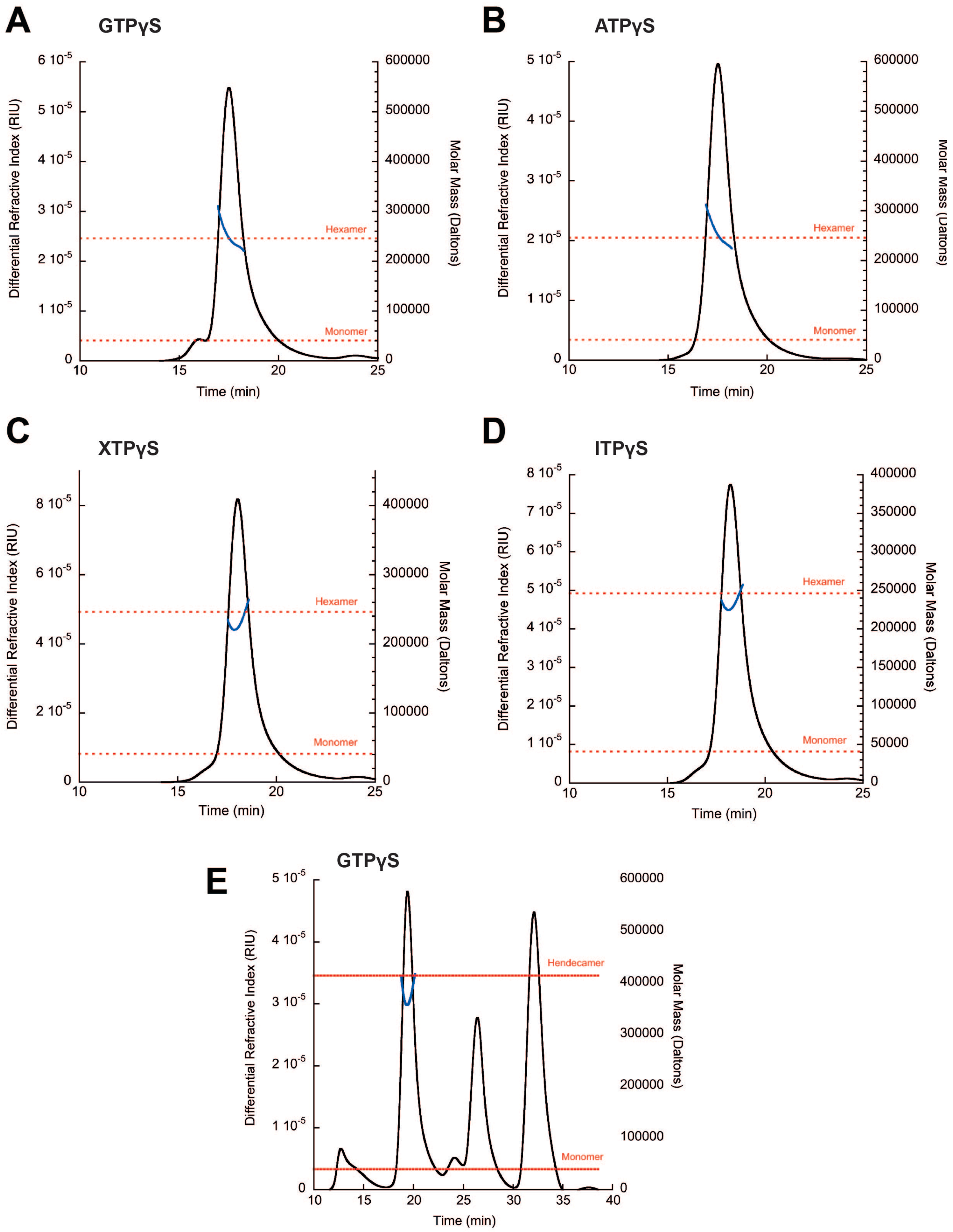


**Figure S4. SEC-MALS analysis of BsuMI.R2 complexes.** A-D, SEC-MALS of BsuMI.R2 in the presence of GTPγS (A), ATPγS (B), XTPγS (C), and ITPγS (D). E. SEC-MALS of BsuMI.R1-BsuMI.R2 complex in the presence of GTPγS.

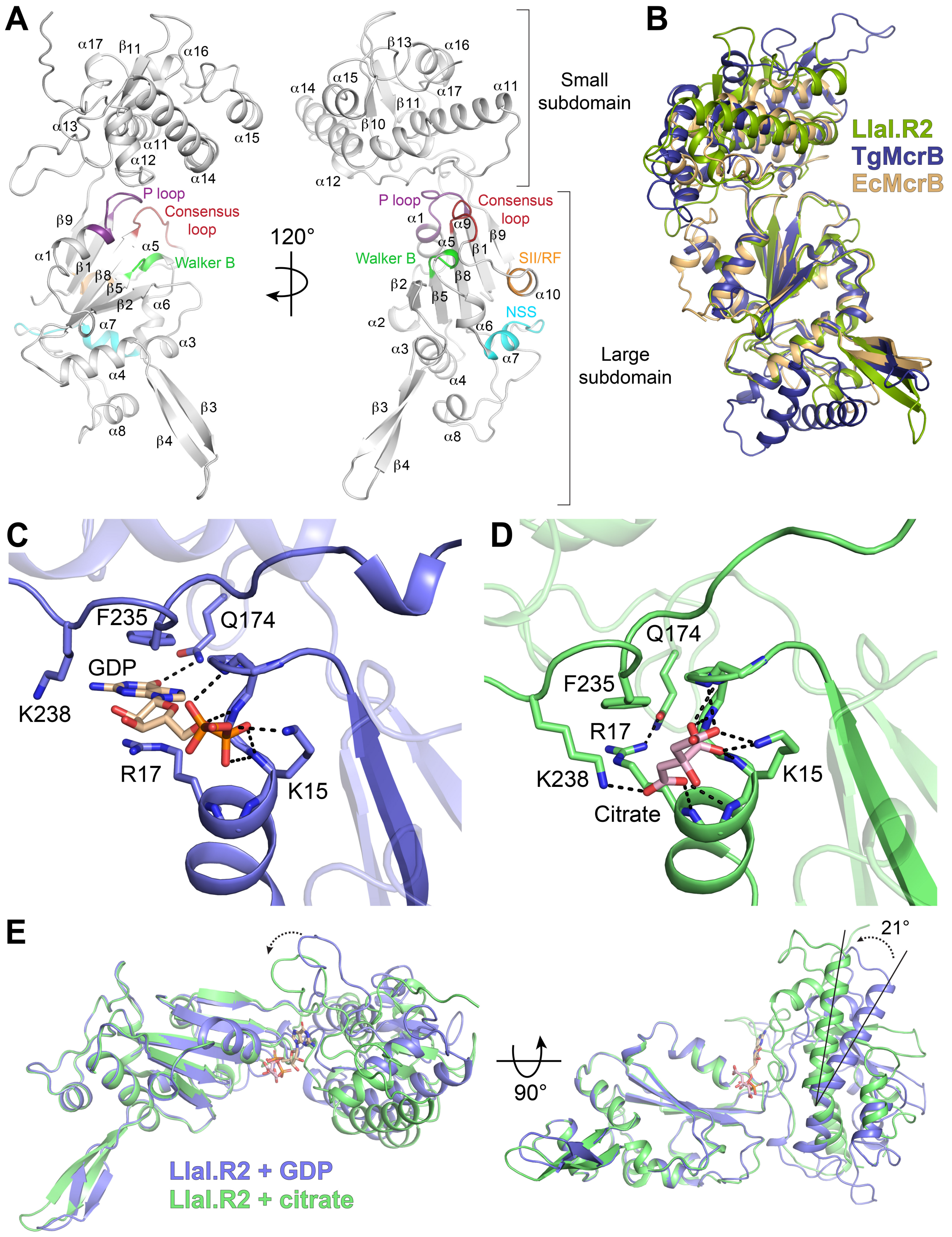


**Figure S5. LlaIR.2 AAA+ fold and comparison of GDP- and citrate-bound LlaI.R2 monomers.** A. Structure of LlaI.R2 monomer. Conserved AAA+ motifs are labeled (see Figure 4A and Supplementary Figure S5 for additional information). B. Superposition of LlaI.R2 (smudged green) with the AAA+ domains from TgMcrB (deep blue, PDB: 6UT5) and EcMcrB (light orange, PDB: 6UT6). C-D. Active site hydrogen bond interactions (dashed black lines) that stabilize the bound GDP (C) and citrate (D), respectively. E. Superposition of GDP- (slate)

and citrate-bound (green) LlaI.R2 monomers in two orientations. GDP and citrate are colored wheat and light pink, respectively.

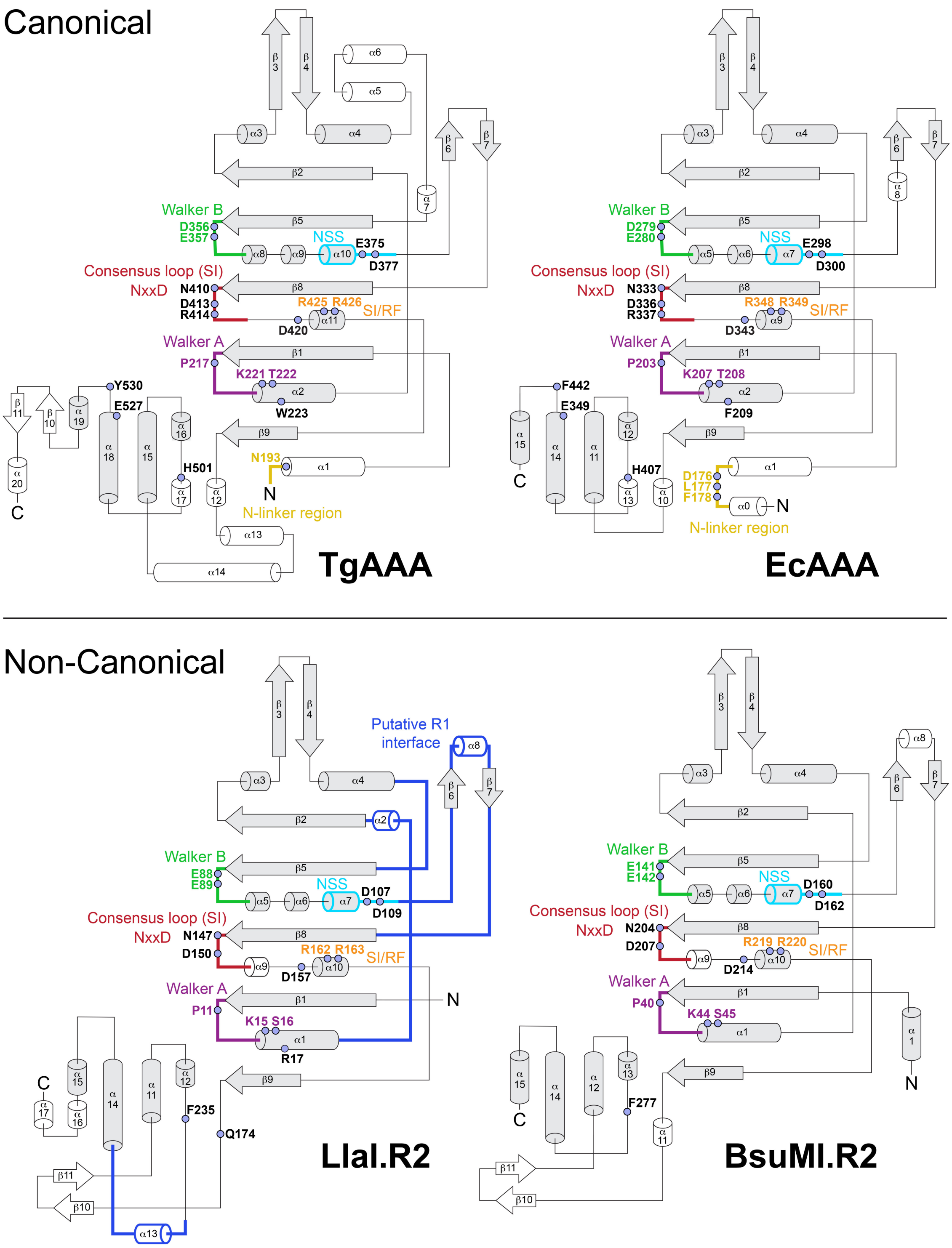


**Figure S6. Topology diagrams of the AAA+ domains from canonical and non-canonical McrB homologs.** Gray shading denotes core fold common to all homologs. Conserved motifs are colored as follows: P loop/Walker A, purple; Walker B, green; nucleotide sandwiching segment (NSS), cyan; McrB consensus loop (Sensor I), red; Sensor II/Arginine finger (SII/RF), orange; N-linker region, yellow. Residues important for nucleotide binding and/or catalytic functions are labeled. Segments contributing to the putative R1-R2 interface are colored blue in the LlaI.R2 diagram. The TcMcrB, EcMcrB, and LlaI.R2 AAA+ topologies shown here were mapped from experimentally determined cryo-EM (PDB: 6UT6, 6UT5) and crystal structures (Figure 2, Supplementary Figure 5A) whereas the BsuMI.R2 AAA+ topology derives from an AlphaFold (9) model.

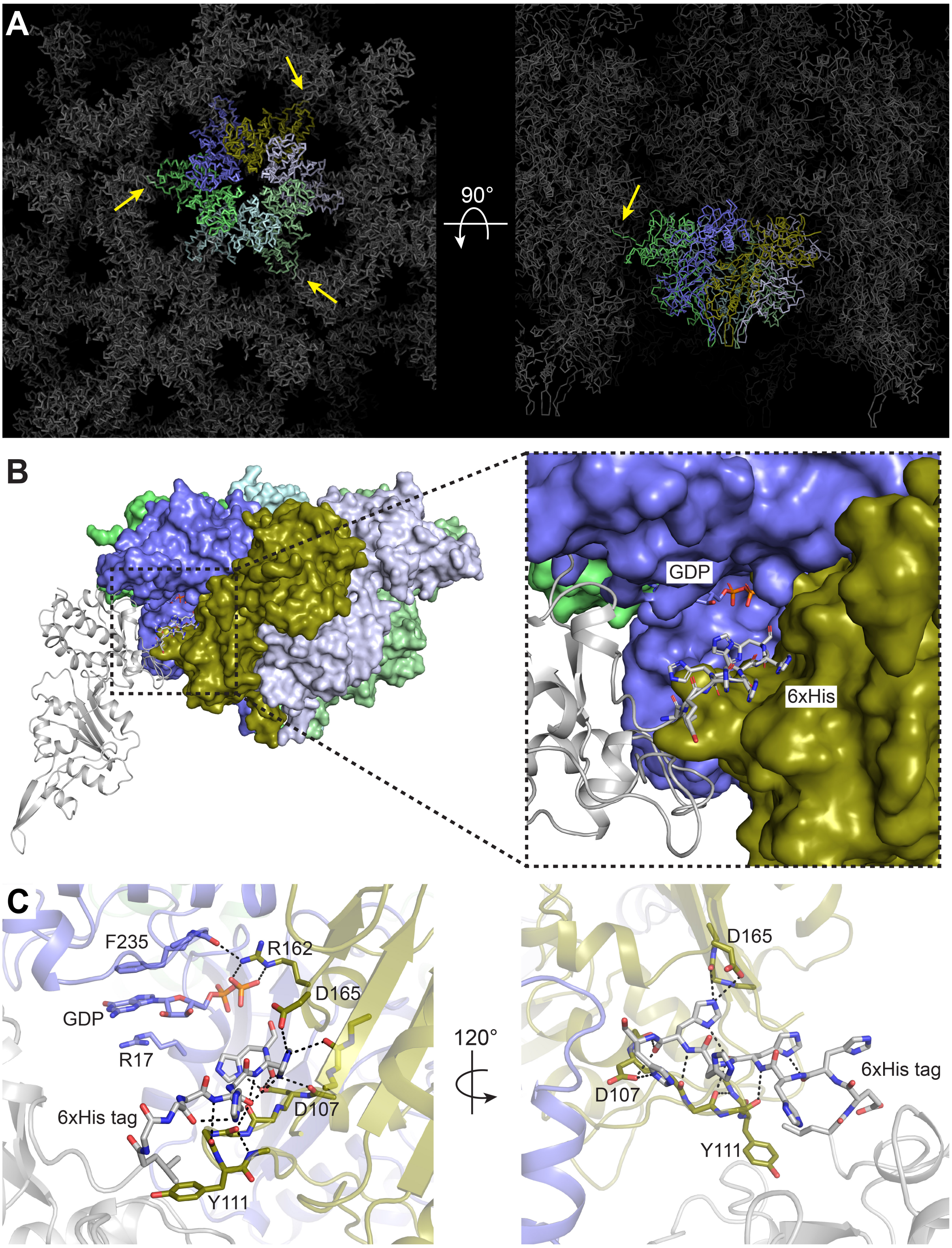


**Figure S7. LlaI.R2 crystal packing and 6xHis tag interactions.** A. LlaI.R2 forms hexamers in the crystal lattice. Yellow arrows denote crystal contacts formed by the C-terminal 6xHis tag wedging into the active sites of neighboring symmetry-related LlaI.R2 monomers. B. Surface view of crystallographic LlaI.R2 hexamer with a symmetry related LlaI.R2 molecule (gray)

rendered as a cartoon. Zoomed insert shows position of the 6xHis tag (gray) relative to the bound GDP in the composite active site between LlaI.R2 monomers within the hexamer (slate and olive). C. The inserted 6xHis tag (gray) forms stabilizing H-bonding interactions (dashed black lines) with the nucleotide sandwiching segment (NSS) that lies between $\alpha$7 and $\beta$6, thereby blocking its interaction with the guanine nucleotide.

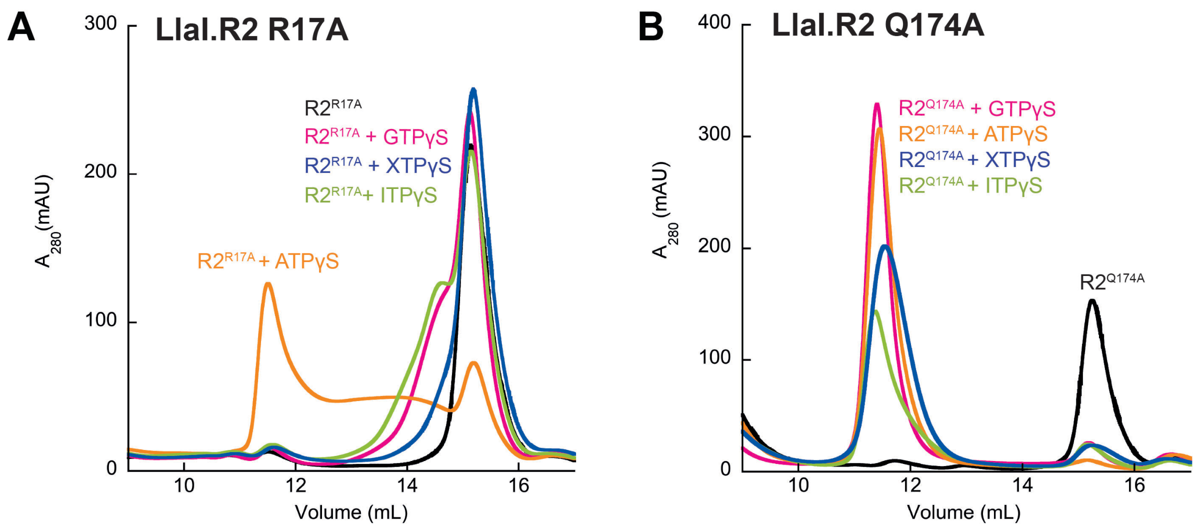


**Figure S8. SEC analysis of nucleotide-dependent oligomerization mutant LlaI.R2 proteins.** Mutant R2 proteins (5 mg/ml) were incubated with nucleotide analogs (2 mM) for 30 minutes at 25°C and then injected onto a Superdex 200 Increase 10/300 GL column. A-B Nucleotide-dependent oligomerization of LlaI.R2 R17A (A) and LlaI.R2 Q174A (B) mutants.

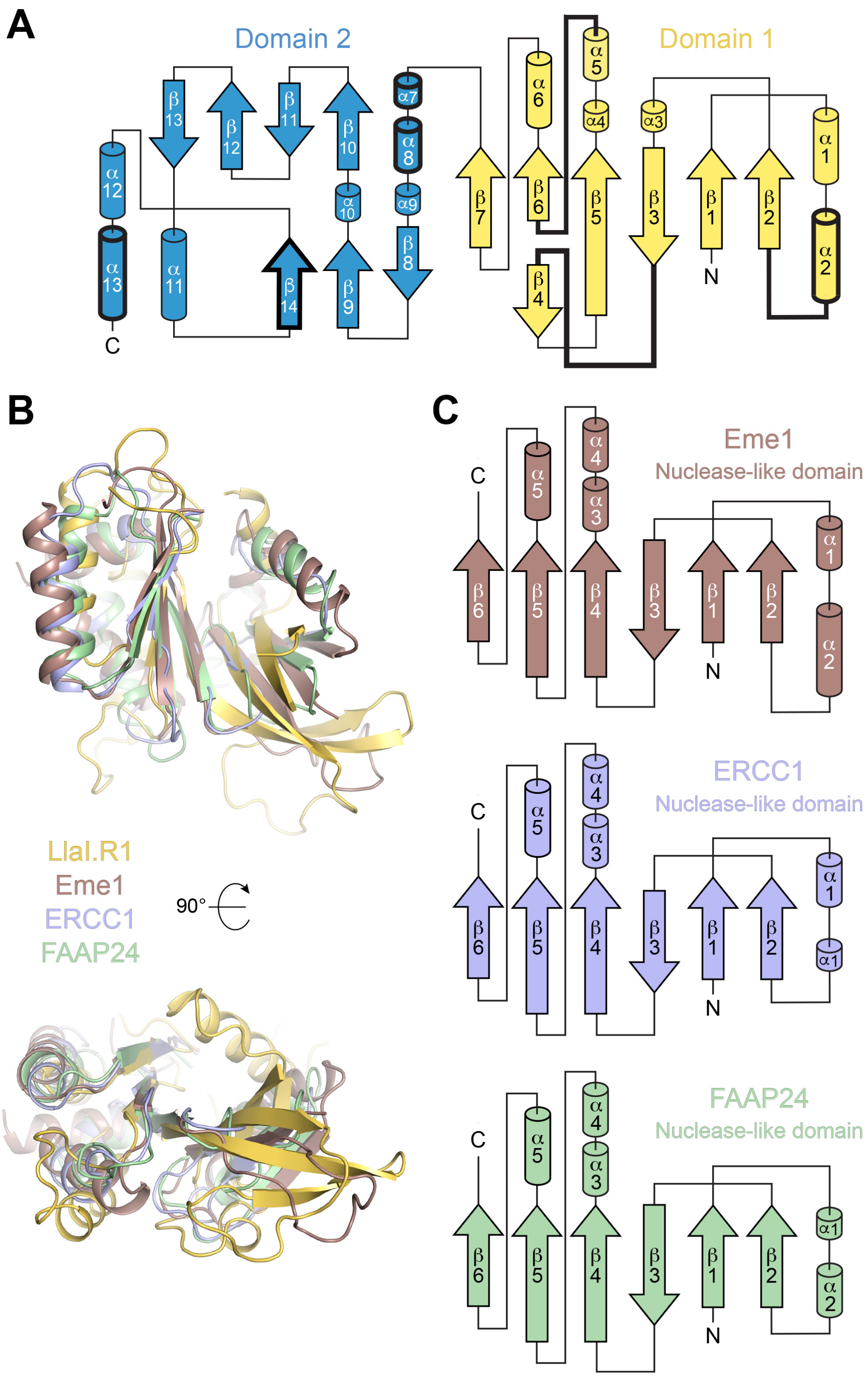


**Figure S9. LlaI.R1 shares structural homology with eukaryotic genome maintenance proteins.** A. Topology diagram of LlaI.R1. Domains 1 and 2 are colored yellow and sky blue, respectively. Thick black lines denote structural segments that contribute to the putative R1-R2 interface. B. Superposition of domain 1 from LlaI.R1 (yellow) with Eme1 (ruby; PDB: 4P0Q,

DALI (10) Z score: 5.3, RMSD: 3.3 Å), ERCC1 (light blue; PDB: 6SXA, DALI Z score: 6.5, RMSD: 3.1 Å), and FAAP24 (pale green; PDB: 4BXO, DALI Z score: 6.1, RMSD: 3.0 Å) nuclease-like domains. C. Topology diagrams of structurally conserved nuclease-like domains in B.

```
LlaI.R1        β1           α1              TT        α2                β2
                1           10         20      30        40        50
LlaI.R1 ......MRKLLLQDG........SYQTKFNKID.....VTPPTGVSKLEYIETKLLQEINNKNPRLNILAIEYLD
Crm     ......MRKLLLQDG........NYQMKLEKID.....VTPPPGVLKLQYIEEIIIRAINDKNPRLHVSSIEYVD
Lem     ......MRKLILQDV........KYQNKFNRID.....VIPPSGISKLKHIETKLLRELNNKNPRLHVSNIEYLD
Lim     ...MKRMRKLLLQDG........SYQTKFNKID.....VTPPTGVSKLEYIETKLLQEINNKNPRLNILAIEYLD
Enf     ......MRKLLLQDG........SYQTKFNKID.....VTPPTGISKLEYIEIKLLQEINNKNPRLHVETIEYLD
Mem     .....MERIILLNDR........QFPDAFLLAG.....VVPPAGVMKIPYVEQKIIQAVNTYNPKLQVQKIEYAA
Pet     .....MKRIILLN.E........SFPNEFKSVN.....IELELEYEKINYIENQIINSVNNYNSKLNVEHIEYLD
Vin     .....MERTLLLESS........GFPEKLELID.....IETPENLLKQSYVEQQVLKSINTYNPKLKVKKVEYIN
Lsm     ...MRTLRKLLLPAS........GFPEKINAIE....SVRIPPTAISKQDFVETAILQAINTYNPKLNLQEVEYSN
Ltm     MKKVRTLRKLLLPAS........GFPEKINAIE....SVRIPPTAISKQDFVETAILQAINTYNPKLNLQEVEYSN
Cam     ......MRILNLNNSLQ...NGSRLPEKFLNIND...STLLGSTPDEIISAVEAKIMKKINSFNNKLNISKIEYGN
Ana     .....MERILYLNDK........TFPDLFYKHP........IYKNNNSFNIENRIVDKINEYNFKLNIKSIEYGD
Sts     .....MERVVLLNSN........AFPSLFNKIVD.....VELVGEELQNFTENKIIQVLNQSNPKINVKSIEYVD
Sth     .....MQRTILLNEK........VFPERYKKIKN.....LSIKGKALNTFIENLIIFKLNEVNNKLNVHKVEYVD
Stp     .....MKRVLYLNKKDGENSSKNAHPQKFYALN.........GIDSSNIFQFEKKILNRINEYNYKLEIEELRYED
Mam     ....MSETKIYLDIRVKNTLIDKKHDKKTGQMIDNNHKRVMVENGIDNLDQYNNIVIERVNEYNPKLNITGIEYVE
Apk     .....MSSVIFLSNKLKESWEKNVMDKKTNEISTNSNLLALRKVRINTIEDYCKLLIEEMNKINSKYMISDVEFVD
Stg     .....MSTIILLNEKLHNAITLEDKDNKNNDSKKKTHISSLKNIGINEVETYYDLLLKKMNEKYPKYKINNIKFVR
Lis     .....MGATILLNKKLKEAWIGENK.GKNSEIKENNNLKALRKEIINIEEYYQKILEKVNERNSKYNVDSINFVD
Lcl     .....MGSKIVLSKRLQKAWTEDKVDSKSSEVKTNPNTIALRKVGISEINDYYNLLIKKMNDINSKYKVNSIKFVD
Enc     .....MKSILVLHKDLKTSYEKLFSDEKKVIKF........KKHGIYSEDDFYKIIIEKINEINSKYGISKIEFSN
Vea     .....MKGIILFSNKLKEAWLAQEKHKKVLSEV.............GGLEVFCNQIIKEVNFVNSKYNVTEVKFVN
            R2                                                      R46

         α3                TT     β3                  β4  TT       β5          TT
LlaI.R1   60          70       80             90         100         110         120
LlaI.R1 INNEF...TGFIGGTYSALIIDEENIPSIPTS......IIDDSRYFTDAIGNVIRQRIMAYVFTSPATKDEGRNIT
Crm     INNEF...PEFIGGIHSAVIVDEENVQSIPTP......QVDDSRYFTNNSGNFIRQRIMAHVFTSPATKDEGRNIT
Lem     INSEF...PNLIRGAYSALIIDEDQIPSIPAT......QIDDSRYFTDSSSNYFRQKIMAYVFTSPATRNEGRDIT
Lim     INNEF...TGFIGGTYSALIIDEENIPSIPTS......IIDDSRYFTDAIGNVIRQRIMAYVFTSPATKDEGRNIT
Enf     IHKGF...SEFIGGKYSAVIVDEKKIPSMPTT......IKDDSRYFVDSGGNVIHQRIMAYVFTSPATKSEGRNIT
Mem     IEAQF...PYYKKGKANGVLIEEFEIHPARSS.....VYRRNGCYVYTRGTKCMCRQILLYLFVSDA.GEDTRNAF
Pet     ISKQF...PSFKTGKSNGLLIEEFTINPKNKDTY.LRNGCYIYSIENDLDKKYICRKILMYLFISNT.SEKSRTSF
Vin     LRTQF...PQVRVSKFTGLIVEDVEIQKEDET.....RFSNAGYFVWEINNKLFARKIVNYVFLSEY.KEKGRNVL
Lsm     INNQFS.NLPVRGGKLFGAIIEDEVINASSED.....YYRNKGYYIWINNGDTVARRIVTYVFILDL.EMDGRNI.
Ltm     INNQFS.NLPVRGGKLFGAIIEDEVINASSED.....YYRNKGYYIWINNGDTVARRIVTYVFILDL.EMDGRNI.
Cam     QTIIP...GVQRTLYSSYIFDRIKISEDERD.....YYLQRSYIVNVIHDQLYAEKMIAYVLLSNS.IQEGRNTT
Ana     VKATS...GLKDSGKFFGLILDIIDINVVKMD...........LPILKLDKGEFYYKIFGYIFISNN.EEVSRSAL
Sts     FNNISFHNFPSKESKFLGVIVDNQIINEKYYKEIKSKIEKENEEYIHENQGEFFYRKIRSYVFLSHP.NSDGRNTK
Sth     FKQLPFSSFPSEVSENIGLIIEDVNISKKQYQ....KFNKNKEGYVYINQGSYYYRKIKAYLYFTQA.EIKSRNIA
Stp     ISK.LFPKANLPSYLHNGIMVREYEVDSNYKI......INDCEEVLFEKDNKCFLRVIDAYVFTTMT.KNDTRNGY
Mam     TPVAV....IGRNAVYSGLIERIDDD...........................GVKSSLAYIFISYP.EVGSRNAF
Apk     IPVPG....NSKSALNAGRIAKKID...........................NEYYTEAYIFISNP.YLGSRNIF
Stg     DPIPF....IKTSPINAGMIVKQIE...........................KEYIPLAYIFVSMP.KLGSRNIF
Lis     EPLPF....LSKQALNAGKIVKYVK...........................DEEKTLAYVYISNP.SLGSRNIF
Lcl     EPVPA....IGSRAINAGLIEREVG...........................GEKIPLAYVFVSNP.NLGSRNIF
Enc     DPIYF....LNKSFKNAGIIVQKDK...........................EKQKGLAYVFLTTP.TLEARNSF
Vea     IDEIPE...LFYLGSQAAGYIVEER...........................RKYILNAYICITNP.DMGSRNAI

         α4       α5               β6              α6              β7       η1
LlaI.R1      130           140           150           160           170           180           190
LlaI.R1 VAQDIFPRLLDYIDQYINSPSYTYANHPFYYISLMTPSGQLQASLLSNYARLNQLDFEYIELFPTGVNFSK.....
Crm     VSQDIFPRLLEYIEKYIDSPSYTYANHPFYYISLMNSSGGLQASILSNYVKLSQLGFEFIELFPIGVDLSR.....
Lem     VTQDIFPRLLDYIEKHIDSPSYTYANHPFYYISLMNTDGGLPASILSNYAKLNQLEIEFIELFATGVNFSK.....
Lim     VAQDIFPRLLDYIDQYINSPSYTYANHPFYYISLMTPSGELQASLLSNYARLNQLDFEYIELFPTGVNFSK.....
Enf     VTQDIFPHLLDYMEKYIDSPSYTYANHPFYYISLMNTNGGLQPSILSNYARLNQLELEFVELFPTGINFSL.....
Mem     VSQTVFPTLLDYAADHLQSPSYSIANHKFCFINILNKKL.TSKMILRHLAGLCAAGMEYVEVFGKDSVVPGD....
Pet     ISQTIFPTLLDYAEDYLDSPSYSLANHKFCFINIVNKKI.TSQMILRHIVSLYLSGMDYIEVFDNNSFIKNN....
Vin     FSQSIFPAIIDYMGDFLSSPSYTIANHPIYFINIMNKEI.TAQSLLKRLAGIVAVRFEYIEVFAETINPKT.....
Lsm     VAQSIFPELIELMDMTIGGPCYKVANHPVYILNMLGSRI.TQSSIYYPIAGMIAADMEYIEVFANTPELVR.....
Ltm     VAQSIFPELIELMDMTIGGPCYKVANHPVYILNMLGSRI.TQSSIYYPIAGMIAADMEYIEVFANTPELVR.....
Cam     TTQQIFPTLADYISHFLSSPSFGYANLPIYYLNIINETI.TSKVMQRRLIEVEQLGIYTIDVFDKNHELYSN....
Ana     FSQTIFPELITIINDSLDSPNFKVSNKPIYLINLIATEI.KASYLLQPLMKLFGIEIVNIFNEWMDDSV.....
Sts     FSQTIFPTTLIYMNEFLNSPMYSIFNHPAYCINLFNGKL.TQNSILKQIYGFSILGVKFLDLFSNIDKTKK.....
Sth     IAQIIFPAILELMGEFINSPMYSIMNHPAYCINLYNDSI.TADSIKRQIYGLNSLGIKYIDLFAKYNEYST.....
Stp     IAQMLFPTLTELMAKFIDSPNYNLLNHPIYVLNVYDKDL..PKKIQRNIVQMKVMGIHYIQVFKNLNNYLE.....
Mam     GSQHLFPGLSKLVDFYIDSPSYDLANLPIYFVNGSFDNI..TNSMLETVISMKLMGINYIQLHNSNSVTVDL....
Apk     GSQQIFPGLSYLIKNYINSPSYELANHPIYFINGSTDSI..TESMIKTLVSIELMGIRYVQLFKSKKEIIKK....
Stg     GAQQLFPGLSYLVKQFINSPSFTLANLPIYFVNGSTDYV..TESMQETITAINLIDVGYVQLFDEESLPPKL....
Lis     GAQQLFPGLSYLINYYISSPAYEFANLPIYFINGSIDPV..TESMQETIMAMNLMNIRYIQLFDDNKLPDGI....
Lcl     GSQQLFPGLSYLVNHYIGSPSYELANLPIYFVNGSTDNV..TESMQETIMAMSLMGVRYVQVFNEETLPAGV....
Enc     GAQQIFPGLSKLVEYFIDSPNFELTNLPVYFINSTIENM..PASKLETIAAISLMNIRYIQIFEPELVPDTL....
Vea     GAQQLFPALSKLVEKYINSPGYELANLPIYFLYGSKDSM..TDSIKQSIIAMELIGVKCIPLFNKGTFLTEDIRLR
```

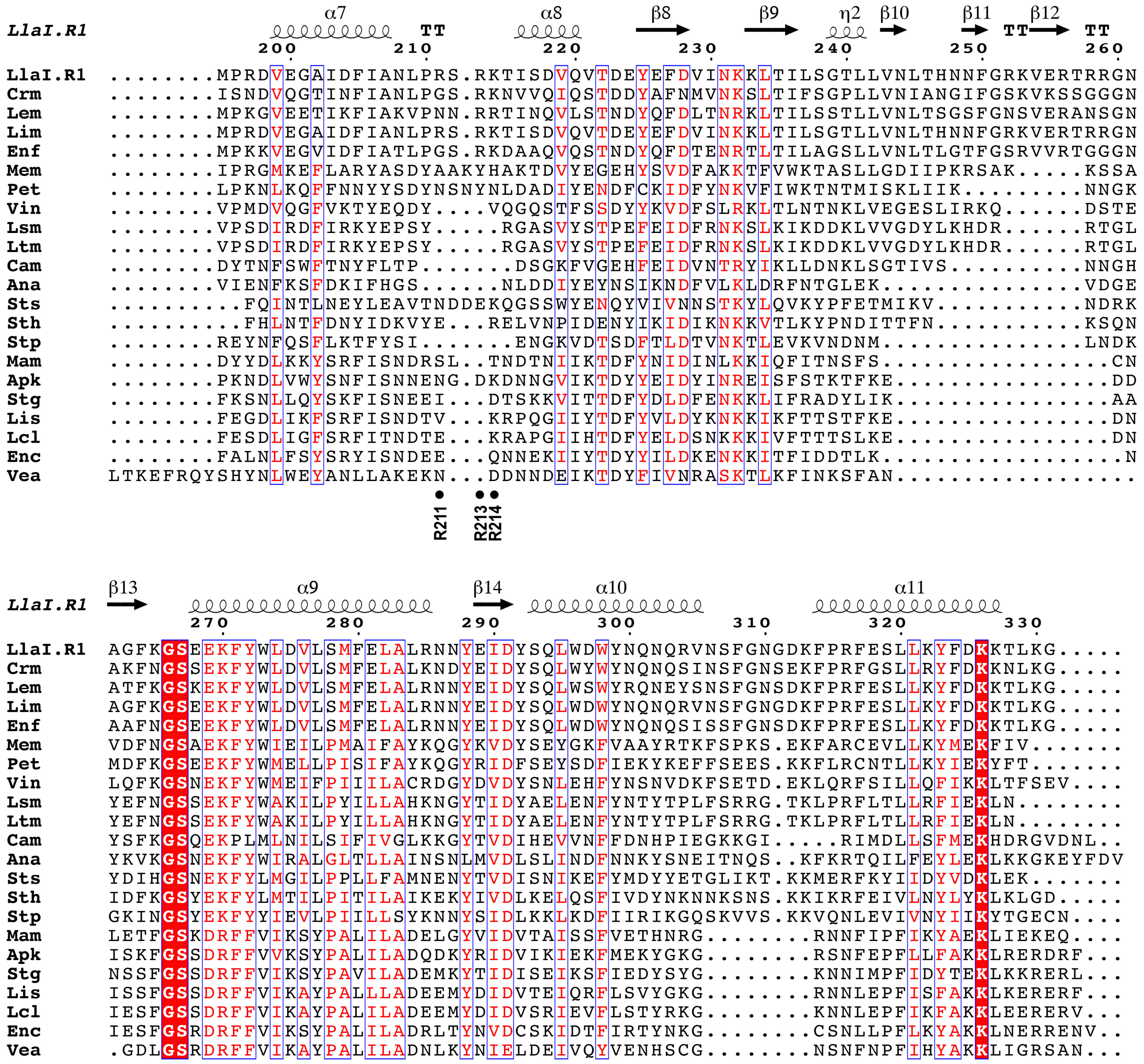

**Figure S10. Sequence alignment of representative LlaI.R1 homologs**. Sequence shading indicates conservation: white text on red background, 100% conserved; boxed red text on white background, 70% conserved. Black circles below the alignment denote residues mutated to alanine to analyze DNA binding (R211A, R213A, and K214A in Patch 1) and the R1-R2 interface (R2 and R46) (See Figures 4-6 and Supplementary Figures S11-S12). Abbreviations for representative homologs are as follows with accompanying NCBI protein accession IDs: Crm, *Carnobacterium maltaromaticum* (WP_229252017); Lem, *Leuconostoc mesenteroides* (WP_223329571); Lim, *Listeria monocytogenes* FLAG-22908 (EAE9683734); Enf, *Enterococcus faecalis* (WP_137685808.1); Mem, *Mediterraneibacter massiliensis* (WP_059066398); Pet, *Peptacetobacter hiranonis* (WP_201415841); Vin, *Virgibacillus natechei* (WP_209464406); Lsm, *Listeria monocytogenes* LM06-01614 (CUK84823); Ltm, *Listeria monocytogenes* PNUSAL010810 (EHT9638667); Cam, *Carnobacterium mobile* (WP_035033457); Ana, *Anaerosphaera aminiphila*

(WP_073185522); Sts, *Staphylococcus saprophyticus* (WP_107643549); Sth, *Staphylococcus haemolyticus* (WP_145429130); Stp, *Staphylococcus pseudintermedius* SPSE-20-VL-NY-GA-0017 (EIA5045348); Mam, *Marinilactibacillus* sp. Marseille-P9653 (WP_225744450); Apk, *Apilactobacillus kunkeei* (WP_187155456); Stg, *Staphylococcus* sp. GDX8P63P (WP_204172600); Lis, Ligilactobacillus salivarius (WP_003699725); Lcl, *Lactococcus lactis* (WP_101916213); Enc, *Enterococcus cecorum* (WP_171312753); Vea, *Veillonella atypica* (WP_005380604).

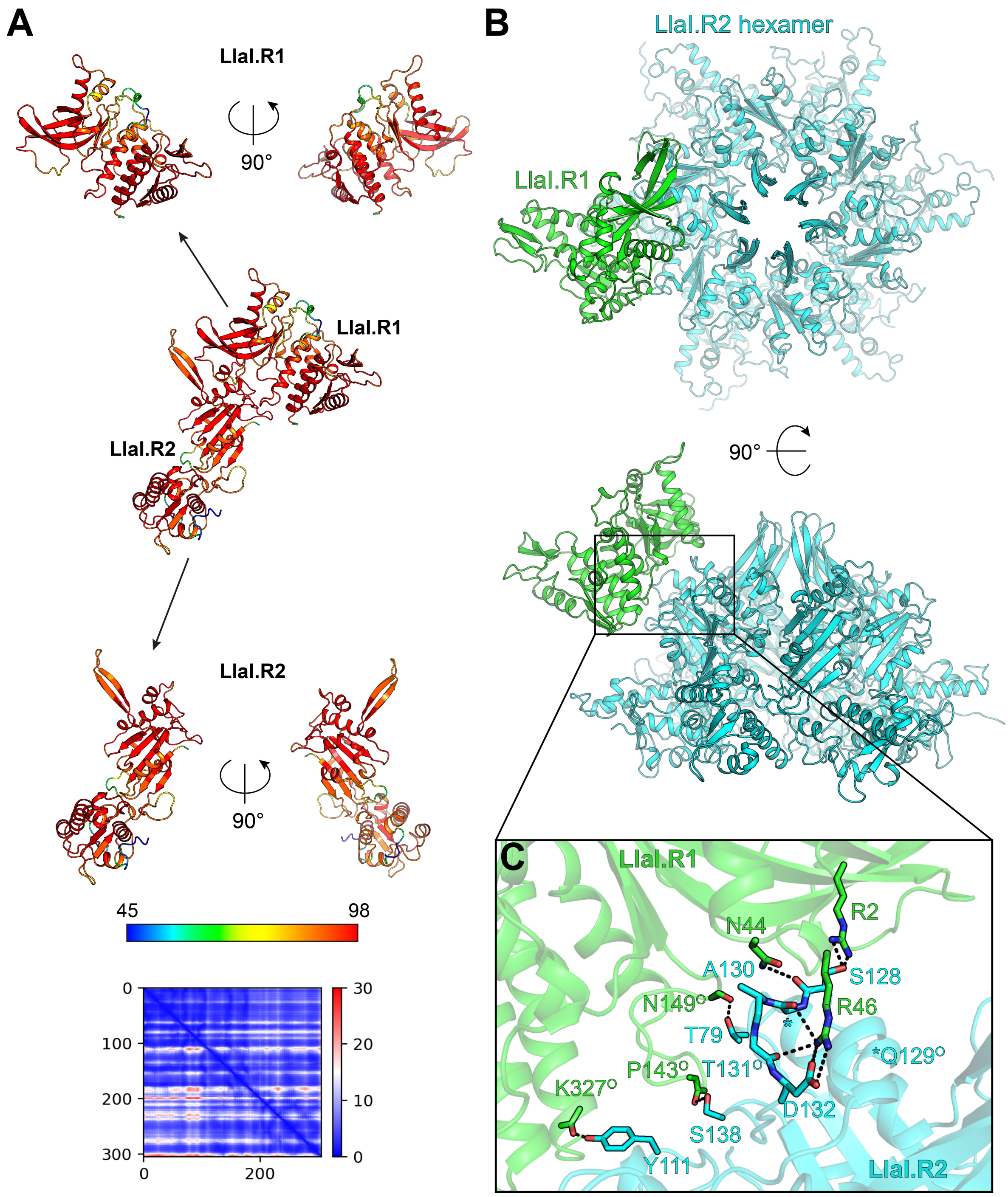


**Figure S11. Structural modelling of LlaI.R1-R2 interactions.** A. AlphaFold (9, 11) generated model of LlaI.R1-R2 complex colored according to the predicted local distance difference test (pLDDT) score (0-100), with values greater than 90 indicating high confidence and values below 50 indicating low confidence. Scale bar denotes per residue confidence coloring for pLDDT score. Individual modelled chains for the R1 and R2 proteins are shown separately for clarity. Predicted aligned error plot for the complex model is shown below. B. Modelled binding of LlaI.R1 (green) to the LlaI.R2 hexmaer (cyan) based on the AlphaFold model in A, depicted in two orthogonal orientations. Black box indicates region highlighted in C. C. Zoomed view of the modelled LlaI.R1-R2 interface. Dashed lines denote predicted hydrogen bonding interactions. "O" superscript signifies backbone carbonyl.

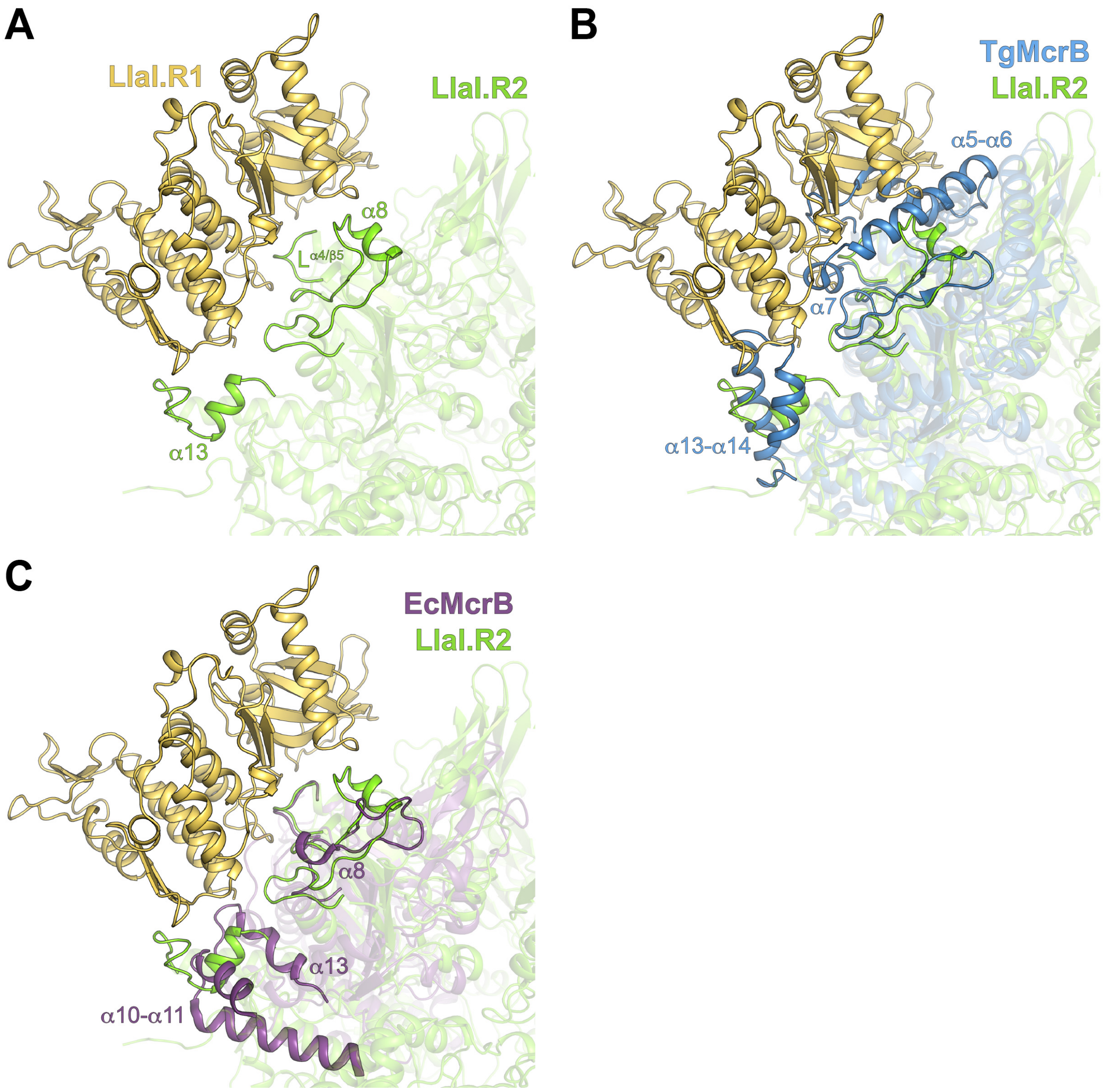


**Figure S12. Evolutionary constraints affecting the association of the DNA-binding and AAA+ modules in McrBC homologs.** A. Structural model (see Figure S11) of LlaI.R1 (light orange) bound to the LlaI.R2 hexamer (green). The positions of the α4-β5 loop ($L^{\alpha4/\beta5}$) and the α8 and α13 helices that contribute of the putative binding interface are highlighted and labeled in LlaI.R2. B, C. Superpositions aligning the TgMcrBC (blue) and EcMcrBC (purple) restriction complexes (PDB: 6UT6, 6UT5) with the LlaIR.2 in the modelled LlaI.R1-R2 complex (A). Structural features that coincide with the putative R1-R2 interface are highlighted and labeled in each assembly. See topology diagrams in Supplementary Figure S6 for further reference.

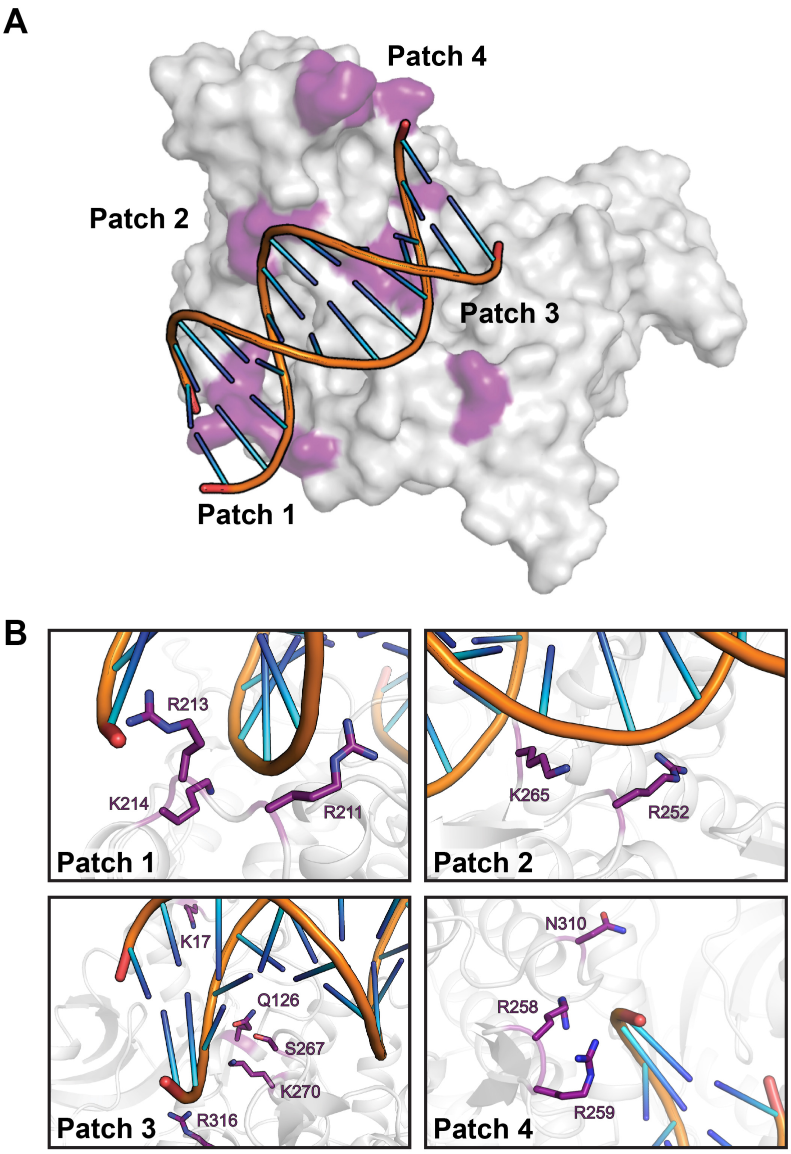


**Figure S13. Structural modelling of LlaI.R1 DNA binding.** A. LlaI.R1 structure (gray) with docked B-form DNA (PDB: 1BNA). Basic residues in patches 1-4 are marked on the surface in purple. B. Zoomed view of putative DNA interactions in basic patches 1-4. Side chains are colored purpled and labelled.

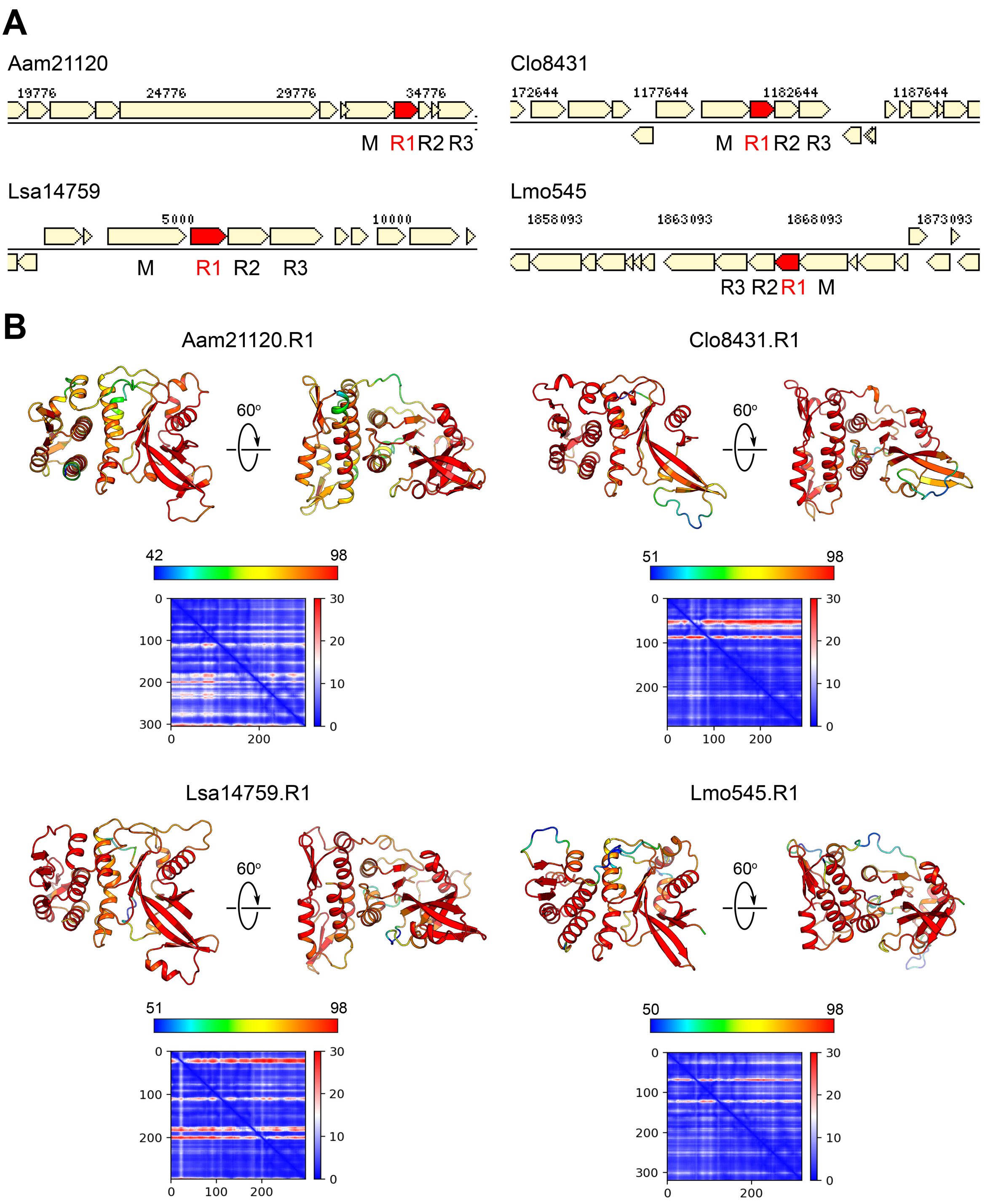


**Figure S14. Structural modelling of LlaI.R1 homologs.** A. Genome neighborhoods of representative LlaI.R1 homologs derived from the DOE IMG/M database (8). R1 genes are marked in red. Species abbreviations are as follows with GenBank (12) and IMG IDs for the associated R1 genes listed in parentheses: Aam21120, *Anaerosphaera aminiphila* DSM 21120 (SHH63106, 2587769317); Clo8431, *Clostridium* sp. DSM 8431 (SFU59092, 2626507161); Lsa14759, *Ligilactobacillus salivarius* BCRC 14759 (ATP37641, 2898119610); Lmo545, *Listeria monocytogenes* PIR00545 (AVU90228, 2900790527). B. AlphaFold (9) models of representative

LlaI.R1 homologs shown in (A). Coloring reflects the predicted local distance difference test (pLDDT) score (0-100), with values greater than 90 indicating high confidence and values below 50 indicating low confidence. Scale bar denotes per residue confidence coloring for pLDDT scores in each model. Predicted aligned error plots are shown below.

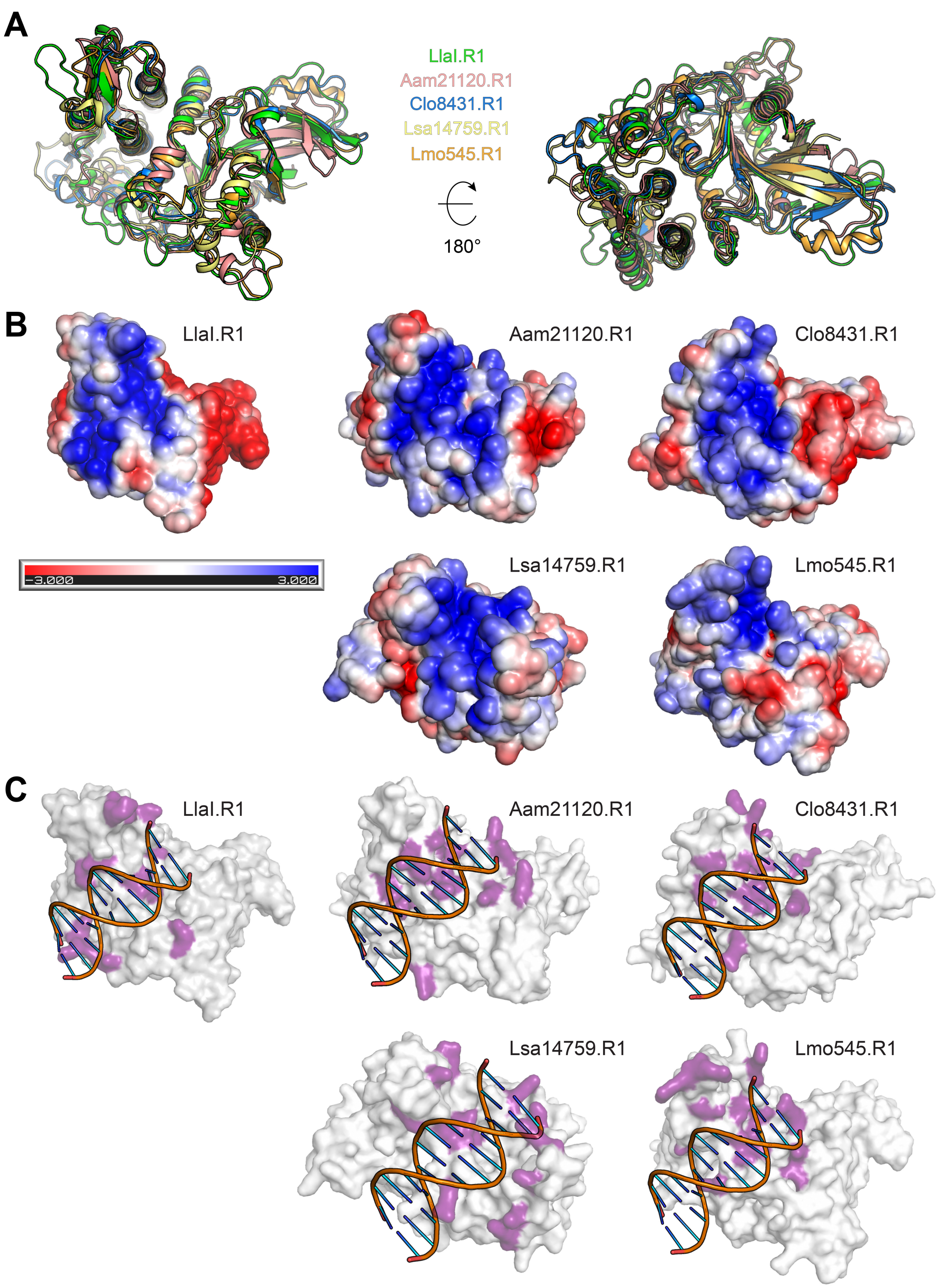


**Figure S15. Structural comparison of LlaI.R1 with modelled R1 homologs identifies a conserved mode of DNA binding.** A. Superposition of LlaI.R1 (green) with AlphaFold (9) models of representative R1 homologs (See Supplementary Figure S14). B. Electrostatic surface across the putative DNA binding face in each of the representative R1 homolog models. Scale bar indicates electrostatic surface coloring from -3 $K_bT/e_c$ to +3 $K_bT/e_c$. E-H. LlaI.R1 (left) is shown in the same orientation for comparison. C. Structural modelling of DNA binding. B-form DNA (PDB: 1BNA) was docked onto each R1 homolog model (gray surface) using HDOCK (13). Side chains predicted to interact with DNA are colored deep purple. The DNA-docked LlaI.R1 structure (left) is shown for comparison.

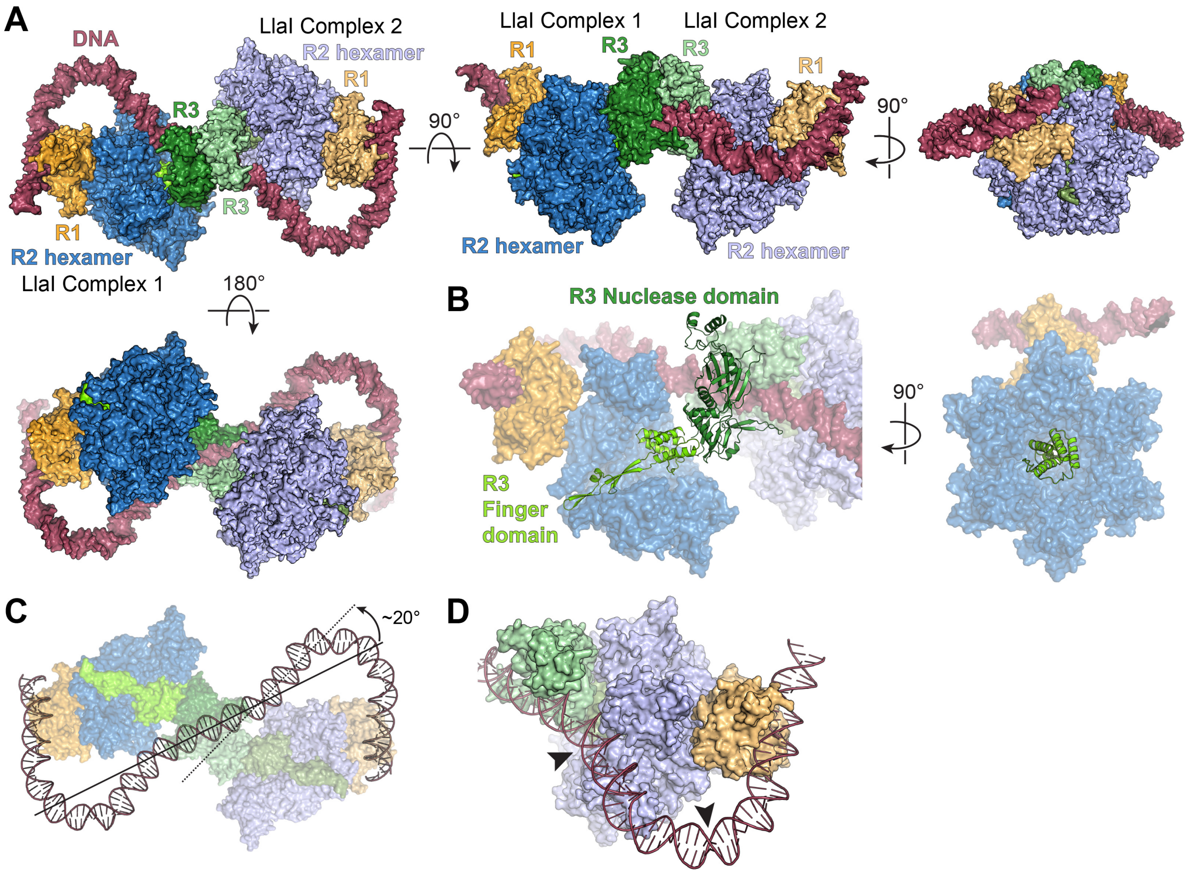


**Figure S16. Structural modelling of assembled LlaI restriction complexes bound to DNA.** A. Top (upper left), side (middle), end (upper right), and bottom (lower left) views of modelled LlaI restriction complexes dimerized on DNA. Each complex is depicted with a stoichiometry of one R1 protein (orange), 6 R2 proteins (blue), and one R3 (green) and shaded in either light or dark colors with DNA colored raspberry. Dimerization via the R3 PD-(D/E)xK nuclease domains is based on interactions observed in the cryo-EM structures of TgMcrBC and EcMcrBC tetradecamer assemblies (PDB: 6UT6, 6UT7, 6HZ5, 6HZ6, 6HZ7, 6HZ8, 6HZ9) (14, 15). Bent DNA substrate was constructed by stitching together segments from the *E. coli* gyrase holocomplex (PDB: 9GVB) and EndoMS (PDB: 5GKE). B. Orientation of the R3 finger (chartreuse) and PD-(D/E)xK nuclease (forest green) domains in the modelled LlaI restriction complex. Two R2 subunits have been removed from the in the side view (left) to highlight the finger domain's show the penetration into the central pore of the hexamer. Right panel is oriented looking down the R2 hexamer pore. C, D. Trajectory of DNA through a modelled LlaI restriction complex dimer. A bend of ~20° would be required to avoid steric clashing with the R2 hexamer as DNA exits the dimerized nuclease domains (C) with additional DNA bending required to bind the R1 subunit (D, black arrows).

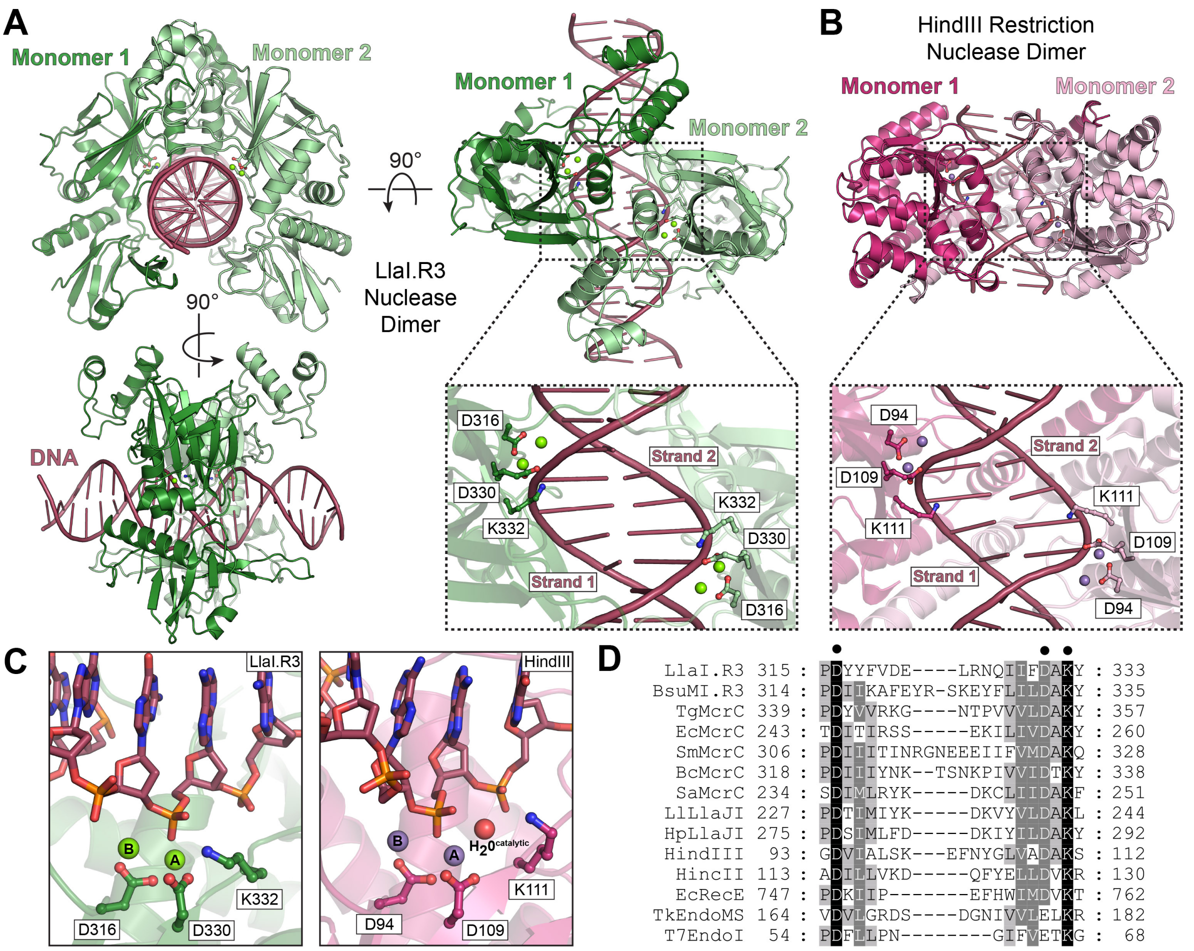


**Figure S17. Modelled LlaI.R3 interactions with DNA.** A. End on (upper left), top down (right), and side (lower left) views of modelled LlaI.R3 PD-(D/E)xK nuclease domains dimerized on DNA (see Supplementary Figure 16 for the orientation of these domains relative to the rest of the LlaI restriction complex). Dashed black box and zoomed inset show how the active sites organize on opposing DNA strands. The putative PD-(D/E)xK catalytic residues that would be critical for metal-dependent DNA cleavage are labelled. Green spheres denote modelled magnesium ions. B. Crystallized structure of the HindIII restriction endonuclease dimerized on DNA (PDB: 3WVH). Dashed black box and zoomed inset show the organization of the active sites on opposing DNA strands. The catalytic residues required for metal-dependent DNA cleavage are labelled. Purple and red spheres mark the positions of bound manganese ions and the catalytic water, respectively, in each active site. C. The spatial organization of catalytic residues and metal ions in the modelled LlaI.R3 active site (left) mirrors that of the crystallized HindIII DNA-bound complex (right), suggesting that McrC homologs a conserved two-metal mechanism for DNA cleavage (16, 17). D. Sequence conservation of PD-(D/E)xK catalytic residues (black circles) across representative McrC homologs, restriction endonucleases, and

DNA replication and repair enzymes. Shading denotes conservation: black boxes with white text, 100% conserved; dark gray boxes with white text, 80% conserved; light gray boxes with black text, 60% conserved. Abbreviations are as follows along with the associated IMG/M (8) or UniProt (18) ID in parentheses: LlaI.R3, *Lactococcus lactis* LlaI.R3 nuclease (UniProt: Q48594); BsuMI.R3, *Bacillus subtilis* 168 BsuMI.R3 nuclease (Uniprot: O34303); TgMcrC, *Thermococcus gammatolerans* EJ3 McrC (IMG/M: 644807739); SmMcrC, *Staphylothermus marinus* F1 McrC (IMG/M: 640109241); BcMcrC, *Bacillus cereus* 03BB102 McrC (IMG/M: 643761467); SaMcrC, Staphylococcus aureus MRSA252 McrC (IMG/M: 637153558); LlLlaJI, *Lactococcus lactis* LlaJI McrC (IMG/M: 642916736); HpLlaJI, *Helicobacter pylori* J99 LlaJI McrC (IMG/M: 637022178); HindIII, *Haemophilus influenzae* Type II restriction enzyme HindIII (UniProt: P43870); HincII, *Haemophilus influenzae* Type II restriction enzyme HincII (Uniprot: P17743); EcRecE, *Escherichia coli* K12 exodeoxyribonuclease 8 (UniProt: P15032); TkEndoMS, *Thermococcus kodakarensis* Endonuclease NucS (UniProt: Q5JER9); T7EndoI, bacteriophage T7 endonuclease I (Uniprot: P00641).
